# Structure-function analysis of 110 phosphorylation sites on the circadian clock protein FRQ identifies clusters determining period length and temperature compensation

**DOI:** 10.1101/2022.09.29.510197

**Authors:** Bin Wang, Elizabeth-Lauren Stevenson, Jay C. Dunlap

## Abstract

In the negative feedback loop driving the *Neurospora* circadian oscillator, the negative element, FREQUENCY (FRQ), inhibits its own expression by promoting phosphorylation of its heterodimeric transcriptional activators, White Collar-1 (WC-1) and WC-2. FRQ itself also undergoes extensive time-of-day-specific phosphorylation with over 100 phosphosites previously documented. Although disrupting individual or certain clusters of phosphorylation sites has been shown to alter circadian period lengths to some extent, how all the phosphorylations on FRQ control its activity is still elusive. In this study, we systematically investigated the role in period determination of all 110 phosphorylated residues reported on FRQ by mutagenetic and luciferase reporter assays. Surprisingly, robust FRQ phosphorylation is still detected even when 84 phosphosites were eliminated altogether; further mutating another 26 phosphoresidues completely abolished FRQ phosphorylation. To identify phosphoresidue(s) on FRQ impacting circadian period length, series of clustered *frq* phosphomutants covering all the 110 phosphosites were generated and examined for period changes. When phosphosites in the N-terminal and middle regions of FRQ were eliminated, longer periods were mostly seen while removal of phosphorylation in the C-terminal tail result in extremely short periods, among the shortest reported. Interestingly, abolishing the 11 phosphosites in the C-terminal tail of FRQ does not only result in an extremely short period, but also causes an over-compensated circadian oscillator under a range of physiological temperatures. When different groups of phosphomutations on FRQ were combined intramolecularly, an additive effect was observed as expected; unexpectedly, arrhythmicity resulting from one cluster *frq* phosphorylation mutants was restored by eliminating phosphorylation at another group of sites, suggesting an epistatic effect between phosphoevents.

**Importance:** Existing in most eukaryotes, circadian clocks are built based on cell-autonomous, auto-regulatory feedback loops in which negative elements feed back to depress their own expression by repressing the positive elements that drive their synthesis. In *Neurospora*, the WCC transcription activator drives expression of FRQ, which complexes with FRH and CK1 to repress the DNA-binding activity of WCC by promoting phosphorylation at a group of residues of WCC. The phosphorylation status of FRQ determines the circadian period length, acting independent of effects of phosphorylation on FRQ half-life. Reflecting this dominant role of phosphorylation, FRQ is subject to substantial phosphorylation at over 100 sites in a time-of-day-specific manner. However, how this plethora of phosphoevents on FRQ controls its activity in a circadian cycle is still elusive, and prior work had shown limited effects of individual phosphosite point mutants. In this study, a series of *frq* mutants targeting multisite phosphorylation within domains of FRQ were generated and analyzed in order to define their roles in period determination. A clear pattern of period altering effects was observed in these *frq* mutants; certain mutants display strong temperature compensation phenotypes, and interestingly, a novel epistatic relationship on rhythmicity between phosphogroups emerged.

## Introduction

Living organisms on earth are persistently under the influence of outside light and dark cycles. To anticipate these environmental changes and better utilize the environmental cues, most organisms have evolved an internal cellular oscillator, the circadian clock, that integrates multiple environmental signals, such as light, temperature, and chemicals, to their internal metabolisms (Dunlap 1996; Dunlap and Loros 2006; Guo and Liu 2010a; Wang *et al*. 2013; Larrondo and Canessa 2018; Diernfellner and Brunner 2020; Zhang *et al*. 2022). Circadian clocks regulate a wide variety of physiological and molecular events in eukaryotes and certain prokaryotes (Michael *et al*. 2015; Karki *et al*. 2020; Ding *et al*. 2021).

Unlike light and chemicals that only function as entrainment cues to the core clock, temperature can impact the core oscillator in several different ways: the period length of circadian clocks remains about the same over a wide range of physiological temperatures—a phenomenon commonly called “temperature compensation” that allows the clock to make accurate time measurements while temperatures undergo large variations in nature; similar to light, both temperature pulses and steps can reset the oscillator, serving as an entrainment factor for clocks (Sweeney and Hastings 1960; Francis and Sargent 1979; Edery *et al*. 1994; Gooch *et al*. 1994; Liu *et al*. 1997); clocks can only oscillate within a limited range of temperatures, outside which the clock will be frozen at a certain phase from which rhythmicity can be resumed if the organism is returned to permissive temperatures (Njus *et al*. 1977).

In *Neurospora, Drosophila*, and mammals, the core oscillator of the clock is built on a transcription-translation negative feedback loop: Negative elements (FREQUENCY [FRQ], PERIODS [PERs], and CRYPTOCHROMES [CRYs]) bring about repression of their transcriptional activators, WC-1 and WC-2 in *Neurospora*, in Clock (Clk) and Cycle (Cyc) in *Drosophila*, and BMAL1/Circadian Locomotor Output Cycles Kaput (CLOCK) in mammals, to terminate their own expression, thereby closing the circadian feedback loop forming the clock (Guo and Liu 2010b; Zhang *et al*. 2011). For example, in *Neurospora*, the White Collar Complex (WCC), a heterodimer comprised of WC-1 and WC-2, serves as the transcriptional activator for the pacemaker gene *frequency* (*frq*) by binding to one of the two DNA elements in the *frq* promoter, the *Clock box* (*C-box*) in the dark (Froehlich *et al*. 2003) or the *Proximal Light-Response Element* (*PLRE*) in the light (Froehlich *et al*. 2002). FRQ interacts with FRQ-Interacting RNA Helicase (FRH) and Casein kinase 1 to repress the transcription activity of WCC by promoting its phosphorylation at a group of residues (Aronson *et al*. 1994c; Lee *et al*. 2000; Cheng *et al*. 2005; Schafmeier *et al*. 2005; He *et al*. 2006; Hong *et al*. 2008; Guo *et al*. 2009; Shi *et al*. 2010; Guo *et al*. 2010; Cha *et al*. 2011; Hurley *et al*. 2013; Lauinger *et al*. 2014; Wang *et al*. 2019).

Protein phosphorylation, as the most common post-translational modification, has been implicated in regulating protein-DNA interaction, protein-protein interaction, protein turnover, enzymatic activity, and subcellular localization, all of which have been shown to control operation of the circadian clocks (e.g., (Luo *et al*. 1998; Diernfellner *et al*. 2009; Lipton *et al*. 2015; Robles *et al*. 2017; Narasimamurthy *et al*. 2018; Luciano *et al*. 2018)). FRQ undergoes dual molecular rhythms in total abundance and phosphorylation (Garceau *et al*. 1997; Liu *et al*. 2000; Ruoff *et al*. 2005). The dynamic phosphorylation status of FRQ is controlled by several kinases and phosphatases including the Casein kinases 1 and 2 (CK1 and CK2), Checkpoint Kinase 2 (PRD-4), Protein kinase A (PKA), Ca/CaM-dependent kinase (CAMK-1), and protein phosphatase 1, 2A, and 4 (PP1, PP2A, and PP4) (Yang *et al*. 2001, 2002; Brunner and Schafmeier 2006; Pregueiro *et al*. 2006; Dunlap 2006; He *et al*. 2006; Huang *et al*. 2007; Cha *et al*. 2008; Diernfellner *et al*. 2019; Wang *et al*. 2021). Newly expressed FRQ is progressively phosphorylated over time and eventually targeted for degradation through the SCF-ubiquitin ligase-recruiting protein FWD-1 (He *et al*. 2003). In *Neurospora*, FRQ is phosphorylated at over 100 residues in a time-of-day-specific manner (Garceau *et al*. 1997; Liu *et al*. 2000; Baker *et al*. 2009; Tang *et al*. 2009) that determine its activities, control its binding partners, and eventually lead to its inactivation (Baker *et al*. 2009; Larrondo *et al*. 2015). Quantitative mass spectrometry data reveal that phosphorylation of distinct regions of FRQ occurs at different phases of the clock, causing opposing effects on its activity and interacting partners over time (Baker *et al*. 2009; Tang *et al*. 2009). *In vitro* kinase assays revealed that CK1 and CK2 account for a large body of FRQ phosphorylation events (Tang *et al*. 2009). In addition to period determination at one temperature, FRQ phosphorylation and related kinases have also been implicated in temperature compensation of the clock at different temperatures (Aronson *et al*. 1994a; Pregueiro *et al*. 2005). For example, CK2 contributes to establishing the temperature compensation of the clock via FRQ phosphorylation at certain residues (Mehra *et al*. 2009). In a recent study, FRQ-CK1 interaction as well as CK1-mediated FRQ phosphorylation have been shown to play a role in regulating the period length across a temperature range (Hu *et al*. 2021). Temperature also controls the ratio of L-FRQ to S-FRQ, derived from different start codons used in translation initiation, which is crucial for maintaining rhythmicity at a low or high temperature (Liu *et al*. 1997; Diernfellner *et al*. 2005; Colot *et al*. 2005) in addition to FRQ phosphorylation.

Mutagenetic analysis of all the plethora of phosphoresidues on FRQ becomes unavoidable in order to fully understand their roles in controlling the pace of the core oscillator. To this end, we engineered and investigated a large number of *frq* phosphomutants covering all the 110 phosphosites for discovering important ones determining FRQ activity, and the large number of FRQ phosphosites was progressively dissected to smaller clusters of phosphoresidues. Taken together, the data show that eliminating certain phosphoclusters in the N-terminal and middle regions of FRQ mainly causes period lengthening while ablation of multisite phosphorylation at the C-terminus results in an extremely short period of 14-15 hrs. Interestingly, impairing phosphorylation of a cluster of residues at the C-terminus of FRQ does not only shorten the period but also leads to an over-compensated clock in a physiological temperature range. Furthermore, unexpectedly, one group of phosphosites on FRQ can be epistatic to another in period determination.

## Results

### A mutagenetic strategy developed to progressively explore roles of the 110 phosphosites on FRQ

A total of 110 phosphosites on FRQ (Figure 1A) have been identified by mass spectrometry (Baker *et al*. 2009; Tang *et al*. 2009), and mutagenetic analyses have been conducted covering only some of these phosphosites. However, how all these phosphoevents regulate FRQ activity has not been systematically studied. In this study, to identify phosphosites on FRQ required for the pace of the circadian oscillator, we adopted a strategy successfully employed in a recent publication by which a small group of phosphoresidues from over 95 sites on WCC was identified for determining the repression of WCC and thereby the closure of the feedback loop (Wang *et al*. 2019). To this end, we engineered a series of *frq* mutants that bear Ala mutations covering all the 110 phosphosites in a group manner (Figure 1A and 1B), and then assayed their role in period determination by tracking bioluminescent signals in real-time.

**Figure 1.**
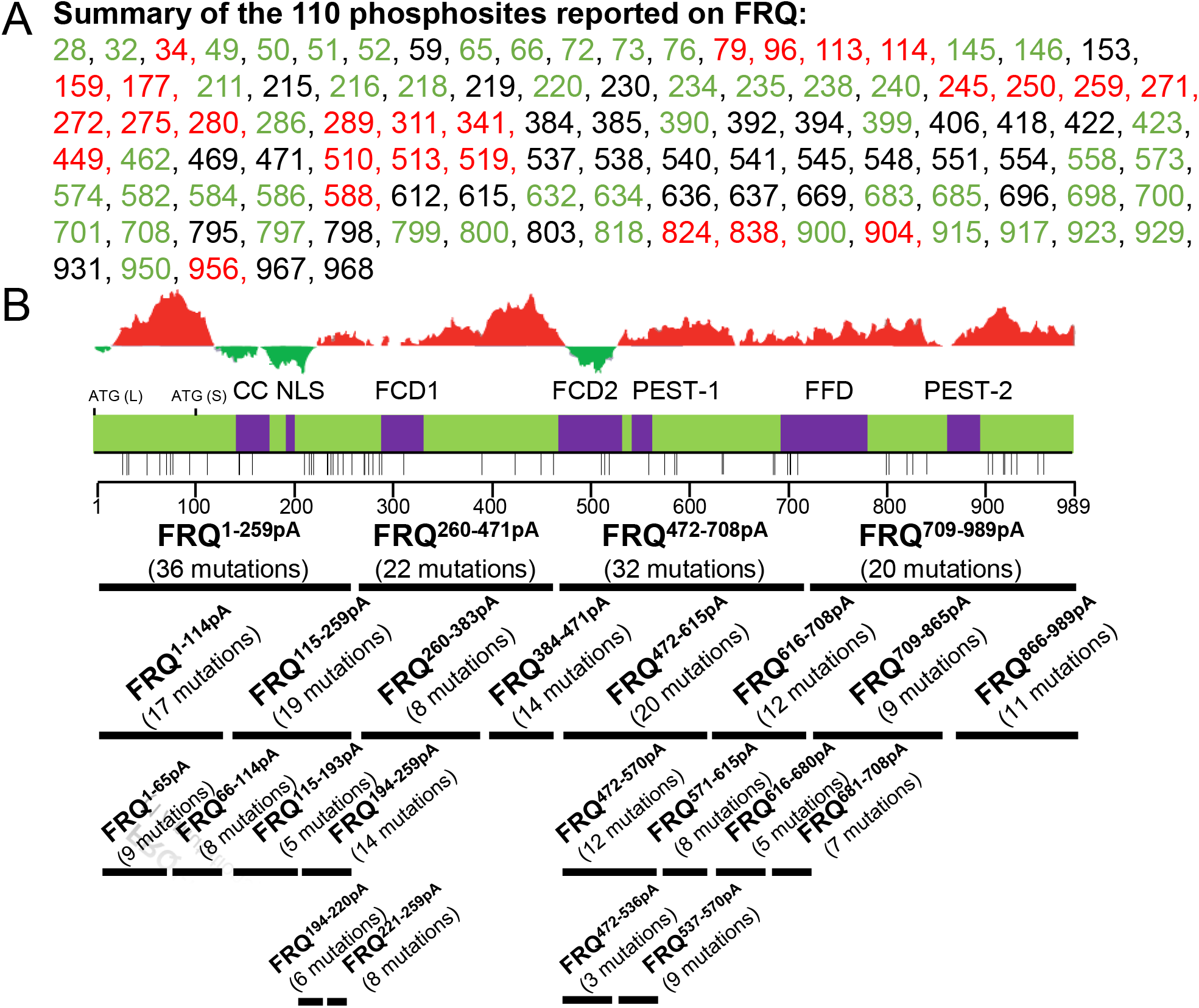
Summary of phosphosites reported on FRQ and *frq* phosphomutants generated in this study (**A**) Summary of the 110 phosphorylation sites from two publications (Baker *et al*. 2009; Tang *et al*. 2009). Numbers represent sites on FRQ at which phosphorylation occurs: Sites reported in (Baker *et al*. 2009), (Tang *et al*. 2009), and both (Baker *et al*. 2009) and (Tang *et al*. 2009) are in black, red, and green, respectively. (**B**) *frq* phosphorylation mutants engineered and investigated in this study. Upper, schematic of FRQ. Each horizontal bar represents a *frq* mutant with phosphosites falling in the region of the bar mutated to Ala altogether while keeping phosphosites outside the region WT. The number of mutations introduced per mutant is in parentheses. ATG (L) is the first start codon used in translation resulting in the full-length FRQ; ATG (S) is an isoform of FRQ translated from the third translational start site (ATG) of the *frq orf*, 99 aa downstream of ATG (L); previously described domains on FRQ are in purple, including the following: CC, coiled-coiled domain; NLS, nuclear localization signal; FCD, FRQ-CK1 interacting domain; PEST-#, pest domains. FFD, FRQ-FRH interacting domain. Each vertical bar below FRQ represents a site phosphorylated by CK1, CK2, or CK1 and CK2 in vitro (Tang *et al*. 2009). Above the diagram is a structural complexity analysis of FRQ amino acids: Red peaks represent disordered regions while green is for structured domains.

### FRQ phosphorylation is detected in *frq^84pA^* but not in *frq^110pA^*

Although over 100 phosphosites have been reported on FRQ, it is still unknown whether this abundance of sites represent all phosphoevents on the protein. To this end, we first engineered two *frq* mutants, *frq^84pA^* and *frq^110pA^* in which the 84 phosphosites reported in (Baker *et al*. 2009) and all the 110 phosphosites from (Baker *et al*. 2009; Tang *et al*. 2009) respectively were mutated to Ala altogether. The clock was assayed in a strain bearing the firefly luciferase gene driven by the *frq C-box* at *his-3* (Larrondo et al., 2015) in which the endogenous *frq* gene was replaced by transformation with the mutant. Luciferase assays reveal that both *frq^84pA^* and *frq^110pA^* are arrhythmic (Figure 2A), suggesting an impaired feedback loop caused by these mutations. The level of FRQ in *frq^84pA^* became extremely low but detectable compared to that in WT (Figure 2B). FRQ phosphorylation in *frq^84pA^* was analyzed using a modified Phos-tag system by which single phosphorylation events on WC-1 and WC-2 could be unambiguously resolved (Wang *et al*. 2019). Surprisingly, despite elimination of all the 84 phosphorylation sites, robust FRQ phosphorylation in *frq^84pA^* is still detected reproducibly by the Phos-tag gel when compared to a lambda phosphatase-treated sample (Figure 2C), suggesting that the 84 phosphosites do not include all major phosphoevents on FRQ. Similar to *frq^84pA^*, the level of FRQ in *frq^110pA^* is dramatically reduced but FRQ phosphorylation in *frq^110pA^* became undetectable when compared to FRQ^110pA^ treated with lambda phosphatase (Figure 2D), suggesting that all major phosphoevents on FRQ have been directly or indirectly eliminated by the 110 mutations introduced.

**Figure 2.**
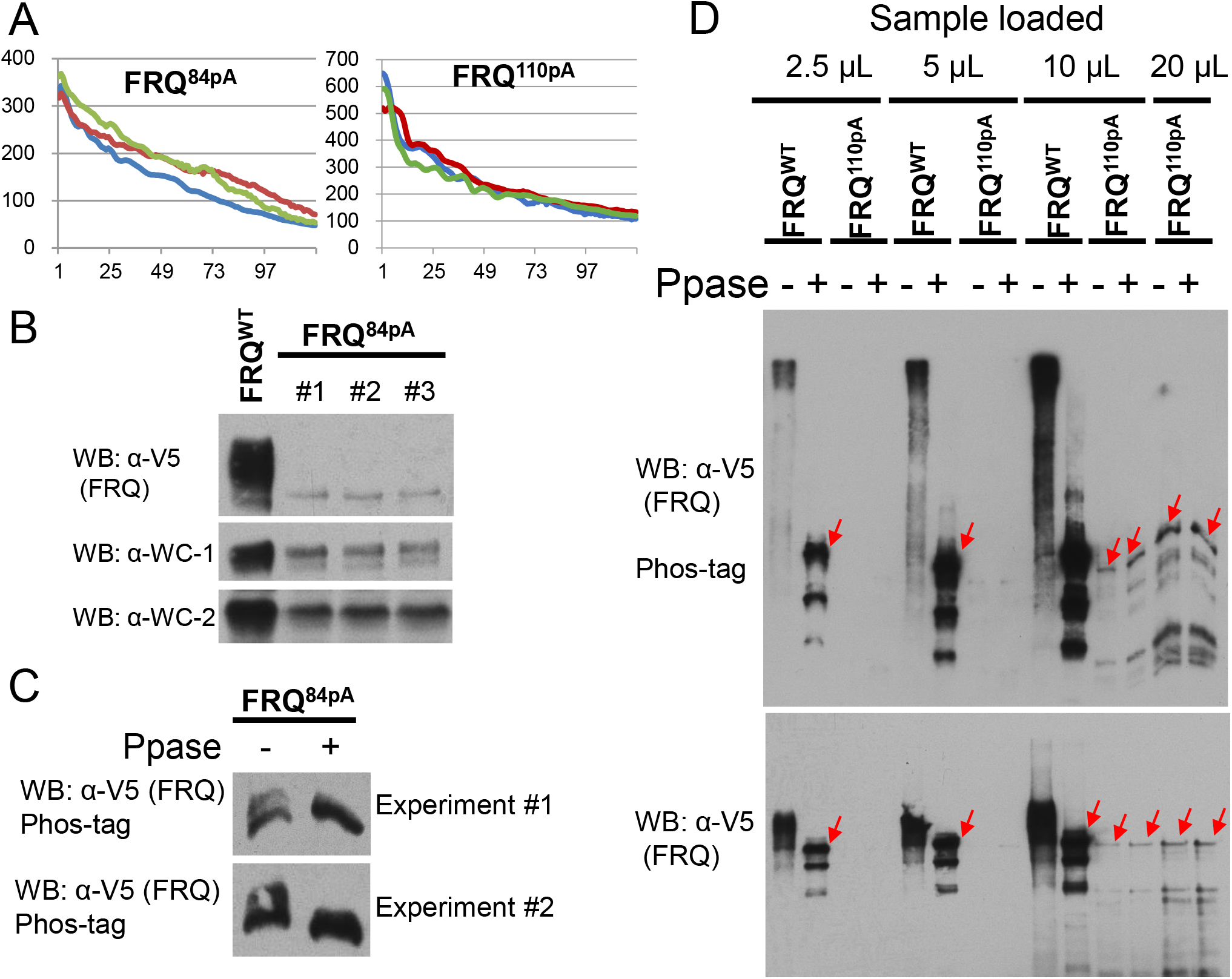
Circadian phenotypes and phosphorylation status of FRQ when all the 84 or 110 phosphosites were eliminated. (**A**) Luciferase assays of *frq^84pA^* and *frq^110pA^* at 25 °C in the dark. *frq^84pA^* and *frq^110pA^* bear Ala mutations to all the 84 phosphosites (reported in (Baker *et al*. 2009)) and all the 110 phosphosites from (Baker *et al*. 2009; Tang *et al*. 2009), respectively (Figure 1A). Strains were synchronized at 25 °C in the light, and after transfer to the dark at the same temperature, bioluminescent signals were recorded by a CCD-camera every hour. (**B**) FRQ, WC-1, and WC-2 expression in WT and *frq^84pA^* by Western blotting. (**C**) Phos-tag gel analysis of FRQ in WT and *frq^84pA^*. FRQ tagged with V5H6 was immunoprecipitated with V5 resin from a constant light culture at 25 °C and then treated with lambda phosphatase to remove phosphorylation. (**D**) similar to (**C**), FRQ in *frq^110pA^* was pulled down with V5 resin from a culture grown in constant light at 25 °C, lambda phosphatase and its buffer supplied by the vendor were added to the washed resin, and the mixture was incubated at 30 °C for removal of phosphorylation. In the gel for Western blot, 2.5, 5, 10, or 20 μL of immunoprecipitated/ phosphatase-treated products were loaded per lane; the upper blot was performed with a regular SDS-PAGE gel, while the lower one was done using a Phos-tag gel. Red arrows point to bands of the full-length FRQ after dephosphorylation, and bands below them are S-FRQ and degradation products of FRQ, which should lack part of the N-terminus because FRQ detected here by WB against V5 is tagged with V5H6 at its C-terminus.

### *frq* mutants identify phosphoresidues affecting period lengths

To directly test the overall effect of FRQ phosphorylation on the period length, we engineered two *frq* mutants, *frq^57pA^* and *frq^27pA^*, together encompassing all the 84 phosphoresidues (Baker *et al*. 2009) mutated to Ala: *frq^57pA^* in which 57 phosphosites falling in amino acids (aas) 1 to 682 of FRQ were mutated to Ala altogether and *frq^27pA^* which bears Ala mutations to the remaining 27 phosphosites found in the region of aa 683-989 of FRQ. Consistent with the arrhythmicity observed in *frq^84pA^* and *frq^110pA^* (Figure 2A), *frq^57pA^* does not develop an oscillating clock while *frq^27pA^* displays a robust rhythm with an extremely decreased period, 14.1 hrs (Figure 3A), shorter than other *frq* mutants bearing point mutations or deletions to the same region of FRQ, such as *frq^S900A^* (19.5 hrs) (Baker *et al*. 2009), *frq^Δ899-989^* (18.7 hrs) (Baker *et al*. 2009) and *frq^C23A^* (18.97 hrs) (Cha *et al*. 2011); this suggests an additive effect played by multiple phosphoevents at the C-terminus of FRQ in controlling the period length. To more specifically test effects of phosphorylations in smaller regions of FRQ, four *frq* mutants derived from *frq^110pA^* were generated, and each of them contains Ala mutants to phosphosites spanning ~200-300 amino acids across FRQ (Figure 1B): *frq^1-259pA^* in which all phosphorylatable residues between aa 1-259 of FRQ were changed to Ala, while keeping aa 260-989 WT (potentially phosphorylatable); *frq^260-471pA^* (phosphosites between 260-471 were changed to Ala); *frq^472-708pA^* (phosphosites between aa 472-708 to Ala); and *frq^709-989pA^* (phosphosites between aa 709-989 to Ala). Luciferase analyses show that *frq^1-259pA^* and *frq^472-708pA^* exhibit a loss of rhythmicity; *frq^260-471pA^* has a lengthened period length (29.4 hrs) while *frq^708-989pA^* displays a decreased period length (14.9 hrs) (Figure 3B), consistent with the phenotype of *frq^27pA^* (Figure 3A). *frq^1-259pA^* bears mutations in and near to the coiled-coil domain that is required for FRQ to interact with itself and other clock components (Cheng *et al*. 2001) as well as mutations near but not within the nuclear localization signal (NLS) (Luo *et al*. 1998), which would seem to explain the lost rhythmicity seen in the mutant; however, that is not the case as shown below for *frq^115-193pA^* and *frq^194-220pA^*. Phosphorylation surrounding the CC and NLS was eliminated in *frq^115-193pA^* and *frq^194-220pA^* respectively, which showed periods of 20.7 and 26 hrs respectively (Figures 5A and 5B), suggesting that abolishing phosphorylation within or near to these domains does not completely impact FRQ function and arrhythmicity in *frq^1-259pA^* is not entirely the result of eliminating phosphorylation within and close to CC and NLS. Because L-FRQ alone is sufficient for maintaining a normal clock at 25 °C, the arrhythmicity of *frq^1-259pA^* should not result from disruption of S-FRQ expression.

**Figure 3.**
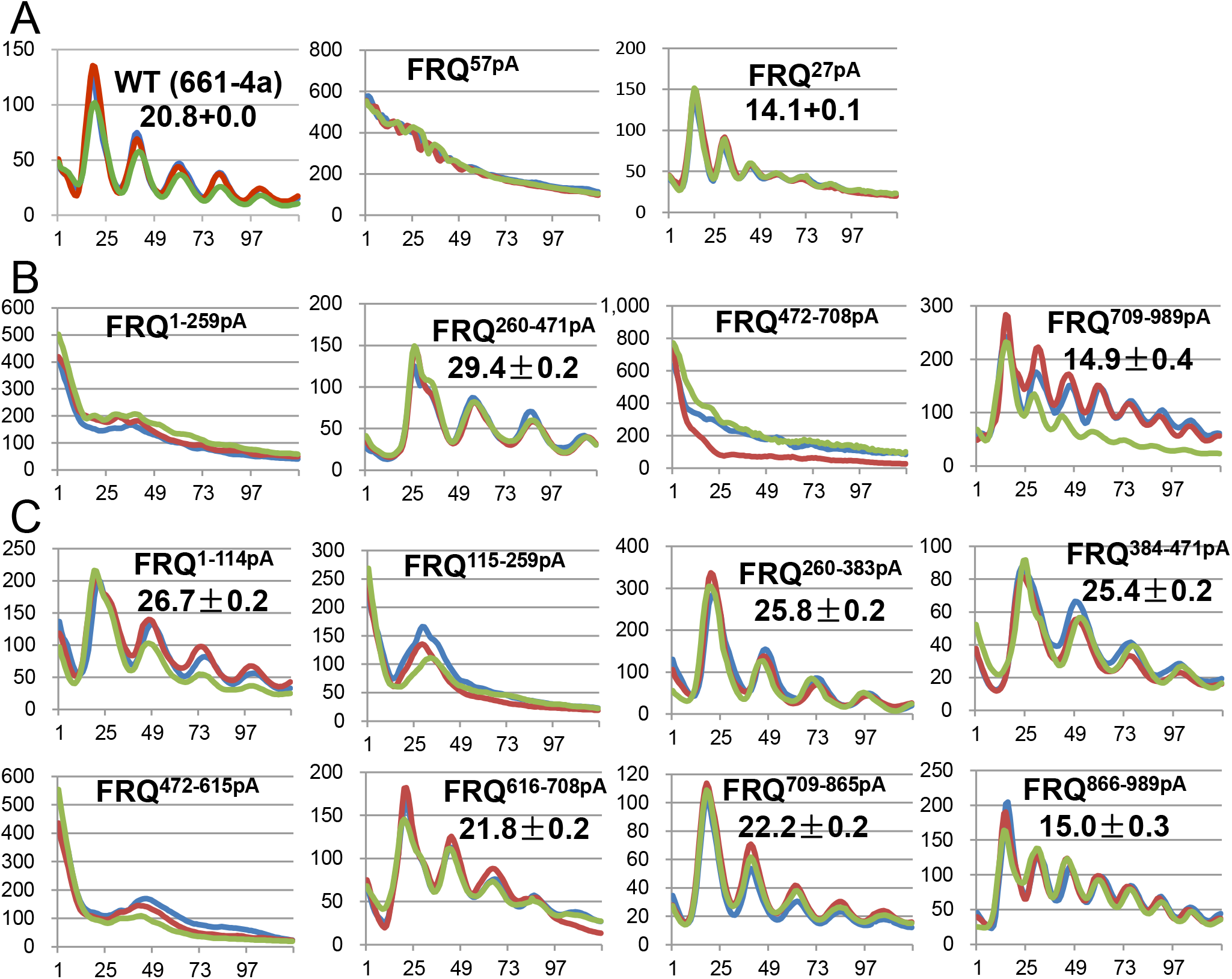
Luciferase analyses of *frq* phosphomutants (**A**) *frq^57pA^* and *frq^27pA^* were analyzed by a luciferase assay at 25 °C in the dark. All the 84 phosphorylation sites on FRQ (Baker *et al*. 2009) were dissected into two *frq* mutants: *frq^57pA^* bearing 57 phosphosites in aa 1-682 mutated to Ala altogether and *frq^27pA^* bearing Ala mutations to the remaining 27 phosphosites. Raw data from three replicates (lines in different colors) were displayed, and time (in hours) and arbitrary units of the signal intensity are on the x-axis and y-axis, respectively. In this and subsequent figures, period length was calculated from three or more biological replicates and is reported as the average +/- the standard error of the mean (SEM). (**B**) Luciferase analyses of *frq^1-259pA^, frq^260-471pA^, frq^472-708pA^*, and *frq^709-989pA^* in the dark at 25 °C. (**C**) Luciferase analyses of *frq^1-114pA^, frq^115-259pA^, frq^260-383pA^, frq^384-471pA^, frq^472-615pA^, frq^616-708pA^, frq^708-865pA^*, and *frq^865-989pA^* in the dark at 25 °C. Strains were cultured in the race tube medium bearing luciferin at 25 °C in the light overnight and transferred to darkness at the same temperature for light production recording by a CCD camera.

To separately follow the impact of these phosphoevents, the 110 phosphorylation sites on FRQ were further divided and mutated into eight additional *frq* mutants (Figure 1B), which were analyzed by real-time luciferase assays as above: consistent with *frq^1-259pA^* and *frq^472-708pA^*, *frq^115-259pA^* and *frq^472-615pA^* are arrhythmic; *frq^1-114pA^, frq^260-383pA^*, and *frq^384-471pA^* show increased period lengths as compared to WT (26.7, 25.8, and 25.4 hrs, respectively); *frq^616-708pA^* and *frq^709-865pA^* have ~WT periods; the period length of *frq^866-989pA^* is 15 hrs (Figure 3C), mostly recapitulating the short period observed in *frq^27pA^* (Figure 3A) and *frq^709-989pA^* (Figure 3B). Expression of FRQ, FRH, WC-1, and WC-2 in all the eight *frq* mutants is comparable to that in WT (Supporting Figure 1); except for *frq^616-708pA^*, the other seven mutants have normal FRQ-FRH interaction; interaction between FRQ/FRH and WC-1/WC-2 is decreased in *frq^384-471pA^, frq^472-615pA^*, and *frq^616-708pA^*, and it becomes undetectable in *frq^115-259pA^* (Supporting Figure 1), consistent with the lost rhythmicity seen in the strain. These data indicate that ablation of certain phosphorylations in the N-terminal and middle regions of FRQ causes period lengthening effects; conversely, removal of phosphorylations at FRQ C-terminus results in an extremely shortened period suggesting an autoinhibitory role for this C-terminal domain. Consistent with the period changes of the *frq* phosphomutants in Figure 3C, canonical *frq* alleles except for *frq^1^* at the N-terminus of FRQ display a lengthened period while *frq^2^* bearing the same mutation as *frq^4^* and *frq^6^* at Ala 895 shows a decreased period (Feldman 1982; Aronson *et al*. 1994a), suggesting that these mutations may impact phosphorylation of other residues, leading to period changes, although they are not phosphorylatable per se.

### *frq^866-989pA^* shows a strong over-compensated clock in a temperature range

Phosphorylation of FRQ and kinases involved, especially CK1 and CK2, have been implicated in controlling temperature compensation (Liu *et al*. 1997; Mehra *et al*. 2009; Hu *et al*. 2021) by which the circadian period length is only slightly altered across a range of physiological temperatures, a conserved mechanism noticed in diverse circadian systems. The eight *frq* phosphomutants in Figure 3C were examined at 20, 25, and 30 °C; *frq^260-383pA^* and *frq^384-471pA^* show a trend similar to that seen in WT; *frq^1-114pA^* and *frq^709-865pA^* display constant period lengths across temperature more so than WT; *frq^115-259pA^* and *frq^472-615pA^* remain arrhythmic, and *frq^616-708pA^* decreased the period with temperature, indicating this strain has an undercompensated clock (Figure 4 and Supporting Figure 2). Interestingly, *frq^866-989pA^* bearing Ala mutations in amino acids 900, 904, 915, 917, 923, 929, 931, 950, 956, 967, and 968 of FRQ demonstrates increased period lengths at higher temperatures, a typical overcompensated clock (Figure 4, bottom left), indicating that phosphorylation of the C-terminal tail of FRQ does not only control the period length but is also involved in maintaining period lengths at enhanced temperatures. Alternatively, because these sites are located close to the PEST-2 domain of FRQ (Gorl *et al*. 2001), whose phosphorylation may indirectly impact the function of the PEST-2 domain, leading to the period change. It is worth noting that the number of mutations introduced to FRQ does not always correlate with the severity of the period alteration. For example, *frq^866-989pA^* bearing 11 mutations displays a dramatically shortened period and an altered pattern of periods across the three temperatures (Figure 3C and Figure 4), while *frq^616-708pA^* with 12 mutations exhibits a WT period at 25 °C and shorter period trend at higher temperatures, and having 9 mutations, *frq^709-865pA^* displays a WT period 25 °C and a constant period at 20, 25, and 30 °C (Figure 3C and Figure 4). *frq^866-989pA^* displays a much stronger period lengthening effect at the higher temperature than *frq^Q2^* which bears Ala mutations to four phosphosites 685, 800, 915, and 929 of FRQ and has a normal temperature compensation phenotype (Mehra *et al*. 2009), suggesting that multisite phosphorylation at the C-terminus of FRQ cooperatively contributes to maintaining the period length at higher temperatures. The translational start codon for S-FRQ is at aa 100, so if arrhythmicity in *frq^1-259pA^* is caused by disruption of the S-FRQ expression, then one would expect to see a similar phenotype in *frq^1-114pA^*; however, it is robustly rhythmic albeit a longer period, excluding the possibility.

**Figure 4.**
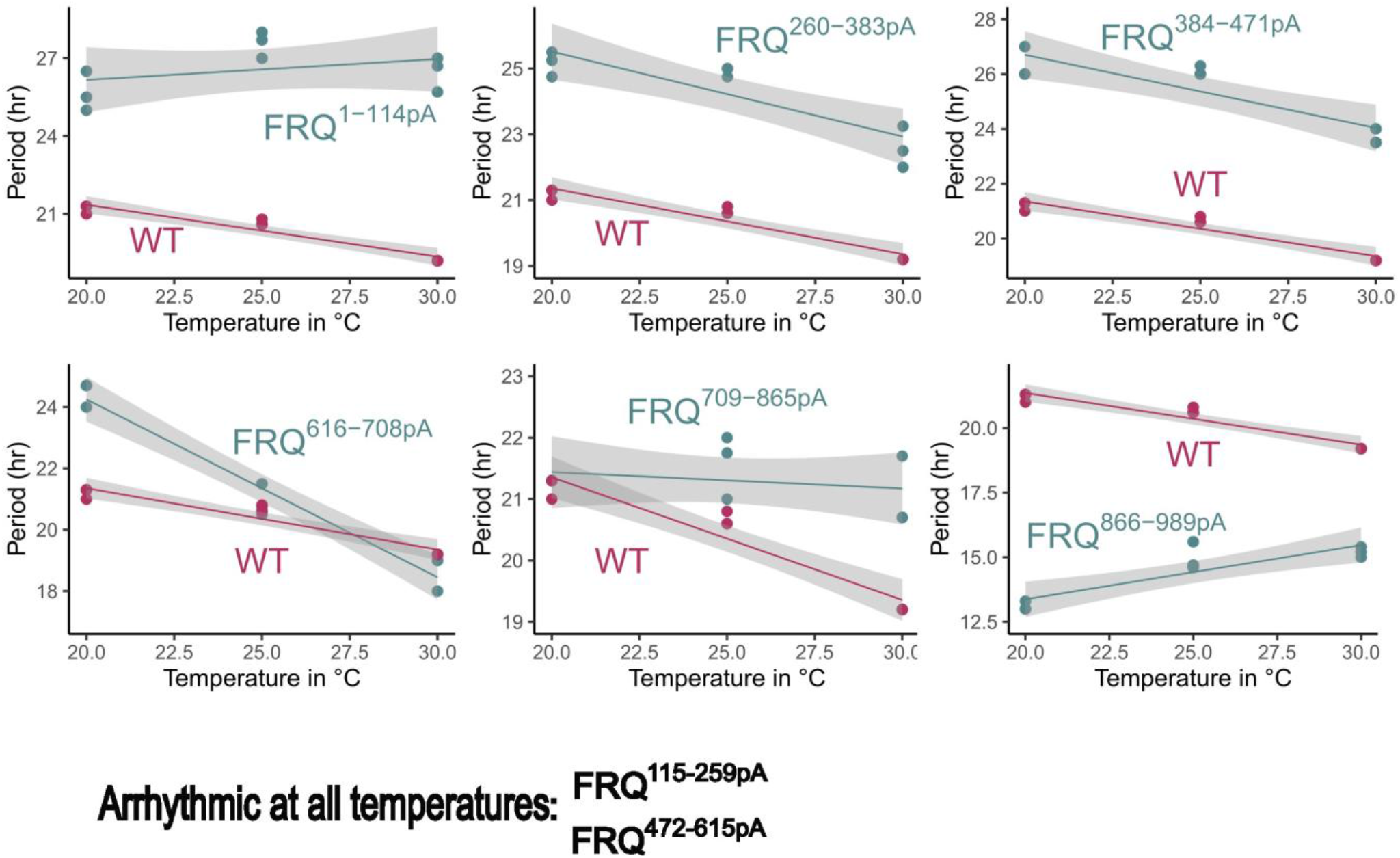
Luciferase analyses of *frq* phosphomutants of *frq^1-114pA^, frq^115-259pA^, frq^260-383pA^, frq^384-471pA^, frq^472-615pA^, frq^616-708pA^, frq^708-865pA^*, and *frq^865-989pA^* at three physiological temperatures, 20, 25, or 30 °C. Strains were synchronized at three temperatures 20, 25, or 30 °C in the presence of light and then transferred to the dark for bioluminescence signal recording at the same temperature used in synchronization. Note: the period length of *frq^616-708pA^* at 30 °C was calculated using the first two cycles only. Temperature in degrees is on the x-axis, and period length in hours is on the y-axis. Raw data are shown in Supporting Figure 2.

### Combination of few key phosphosites, but no single phosphosite, on FRQ is required for temperature compensation of the clock

Given that our mutational analysis of FRQ phosphosites revealed specific domains on FRQ involved in temperature compensation, we investigated at a more detailed level the involvement of single, double, or triple phosphosites on FRQ in temperature compensation. FRQ phosphosite mutants from Baker *et al*. (2009) were crossed to the *C-box-luciferase* reporter and two siblings from each cross were screened at 20, 25, and 30 °C (n=3 at each temperature) (Supporting Table 1). The negative control of *ras-1^bd^* (WT) had normal temperature compensation, and the positive control, *ras-1^bd^, prd-3* (Mehra *et al*. 2009) was overcompensated as expected. Most FRQ phosphosites, when mutated, did not perturb temperature compensation, even when period length was changed (Figure 5A; Supporting Table 1). However, mutation of S5438A & S540A, and mutation of S538A & S540A & S548A on FRQ resulted in overcompensation, i.e. period length increased as temperature increased (Figure 5B). The additional mutation of S548 to alanine increased the period length dramatically, and also caused arrhythmicity at 30 °C, suggesting that this site acts synergistically with the others in this cluster. Mutation of S573A & S574A caused modest undercompensation (Figure 5C). Statistical differences between period length at low versus high temperatures determined using student’s t-test (Figure 5D) indicate that no single phosphosite alone is responsible for period modulation with temperature. Rather, only mutation of a combination of several key phosphosites can perturb temperature compensation, and phosphosites on FRQ that contribute to undercompensation or overcompensation phenotypes are distinct.

**Figure 5.**
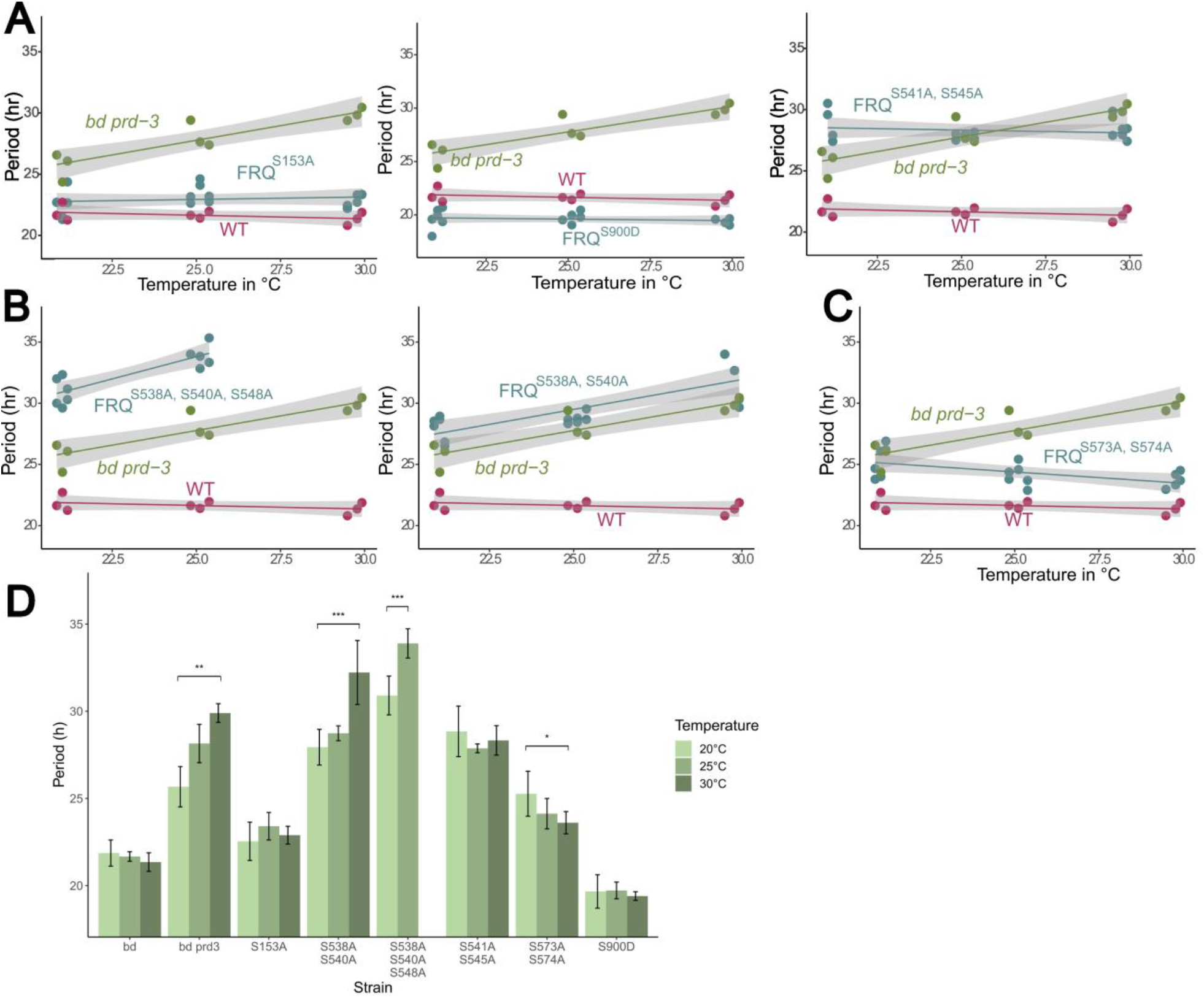
Combination of few key phosphosites on FRQ are required for normal Temperature Compensation. FRQ phosphosite mutants from Baker *et al*. (2009) were screened for temperature compensation defects by crossing in a transcriptional *frq* luciferase reporter into each strain. Strains were entrained on a 12/12 light dark cycle for 2 days at 25 °C, and then transferred to the dark at 20, 25, or 30 °C to record luciferase oscillations. The negative control (labeled WT) was *ras-1^bd^* and the positive control for temperature compensation defect was *ras-1^bd^, prd-3*. (A) Nearly all strains screened showed normal temperature compensation profiles, regardless of period length. Representative examples show WT, long, and short period lengths with normal temperature compensation. (B) Two strains were overcompensated against temperature, FRQ^S538A, S540A^ and FRQ^S538A, S540A, S548A^. FRQ^S538A, S540A, S548A^ was arrhythmic at 30 °C. (C) One strain was undercompensated against temperature FRQ^S573A, S574A^. (D) Period lengths of strains depicted in A, B, and C at each temperature tested. Two siblings from each cross were screened, n=3 at each temperature. Student’s t-test was used to determine statistical significance between period length at 20 °C versus 30 °C (25 °C vs. 30 °C for FRQ^S538A, S540A, S548A^). P-value of * is ≤ 0.05, ** is ≤ 0.01, and *** is ≤ 0.001. Strains without an asterisk were not significant. Supplemental Table 1 lists period lengths for all strains tested, including those not depicted here. Supplemental Figure 3 shows luciferase traces for strains shown in A, B, and C.

### Further defining phosphosites in the arrhythmic mutants of *frq* in Figure 3C

Because eliminating phosphorylation in aa 115-259 or 472-615 resulted in lost rhythmicity (Figure 3C), more *frq* mutants bearing fewer mutations were generated to these and their neighboring regions (Figure 6A) in order to understand the roles of these phosphoevents. *frq^1-65pA^* bearing 9 mutations displays a WT period length, while *frq^66-114pA^* with 8 point mutations shows a long period length similar to that in *frq^1-114pA^*, suggesting that the period effect of phosphorylations in aa 1-114 is mainly caused by phosphorylations in aa 66-114 (Figure 6A). The period of *frq^115-193pA^* is only slightly shorter than WT while *frq^194-259pA^* is still arrhythmic as *frq^115-259pA^* (Figure 6A), indicating the arrhythmicity in *frq^115-259pA^* is mainly caused by losing phosphosites in aa 194-259. It seems that phosphorylation does not impact FRQ dimerization, because *frq^115-193pA^* remains ~WT although it bears mutations close to and within the CC domain (aa 143-176). Although *frq^472-615pA^* is arrhythmic (Figure 3C), *frq^472-570pA^* shows a long period length of 46.3 hrs, which to our knowledge is the longest period length in *frq* phosphomutants to date, and *frq^571-615pA^* shows 26.4 hrs (Figure 6A). *frq^616-680pA^* displays a long period, 26.1 hrs while *frq^681-708pA^* is only slightly shorter (Figure 6A). *frq^616-708pA^* shows an intermediate period between *frq^616-680pA^* and *frq^681-708pA^*, which suggests that the effect of phosphorylations in one region can be averaged by phosphorylations in the neighboring region. Bearing mutations near the FFC domain of FRQ, *frq^616-708pA^* has less FRH complexed with FRQ (Supporting Figure 1) but it still maintains a ~WT period (Figure 3C), suggesting that the negative arm complex in the mutant still functions normally by a decreased level of FRH binding to FRQ.

**Figure 6.**
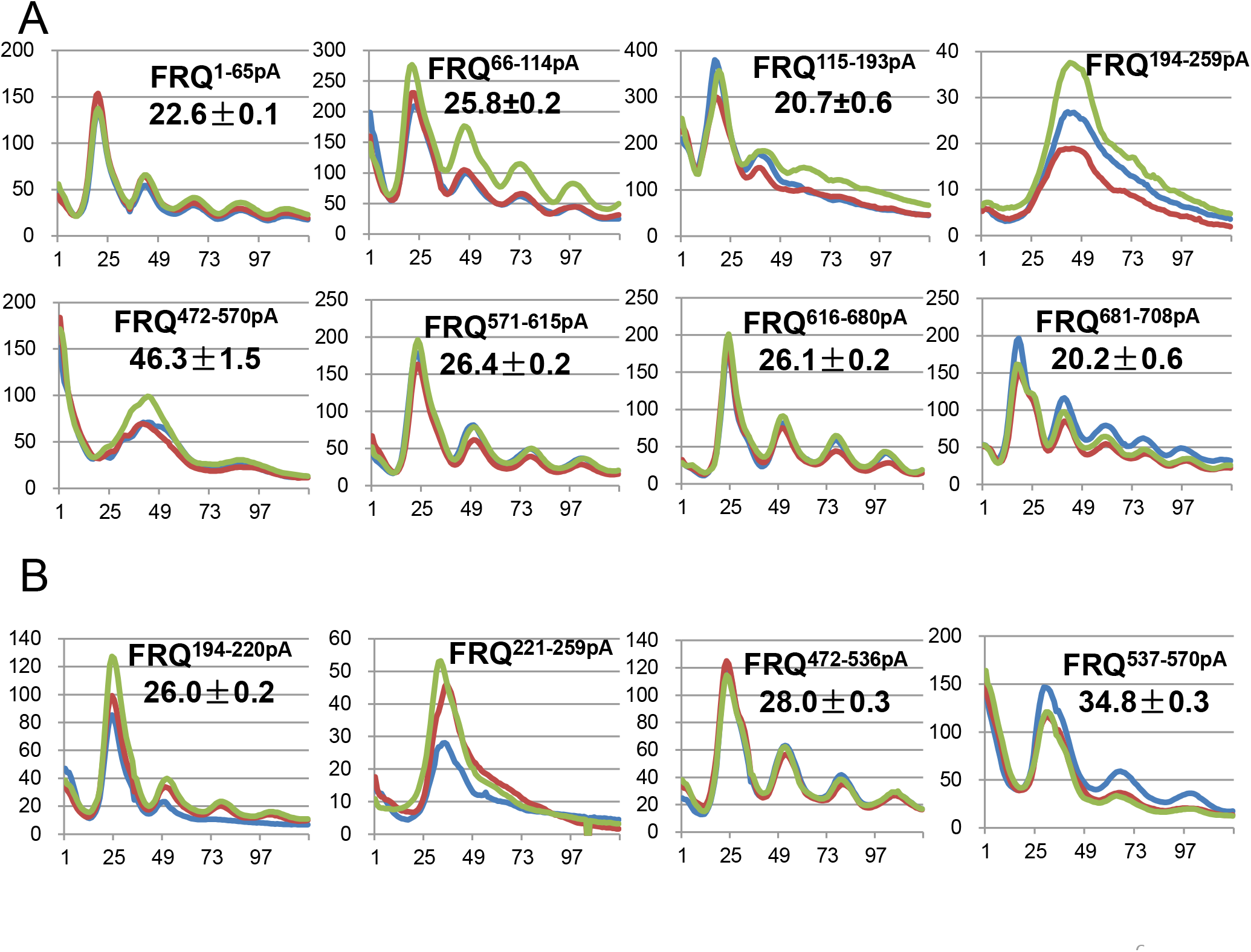
Further dissecting FRQ phosphorylation events falling in amino acids 1-259 and 472-708. **(A)** Luciferase analyses of *frq* phosphomutants, *frq^1-65pA^, frq^66-114pA^, frq^115-193pA^, frq^194-259pA^, frq^472-570pA^*, *frq^571-615pA^, frq^616-680pA^*, and *frq^681-708pA^* at 25 °C. Note: the period length of *frq^472-570pA^* was calculated only from two available circadian cycles. (**B**) Luciferase analyses of *frq^194-220pA^, frq^221-259pA^*, *frq^472-536pA^*, and *frq^537-570pA^* at 25 °C.

To understand why loss of phosphorylation between aa 194-259 causes arrhythmicity (Figure 6A), two additional mutants *frq^194-220pA^* and *frq^221-259pA^* were generated and assayed by luciferase analyses: *frq^194-220pA^* is 26 hrs and *frq^221-259pA^* is arrhythmic (Figure 6B). *frq^194-220pA^* has mutations to phosphosites near the NLS (aa 194-199) but it is still robustly rhythmic albeit a long period, suggesting that phosphorylation does not control nuclear localization of FRQ, which is required for its function in the clock (Luo *et al*. 1998). The arrhythmicity seen in *frq^221-259pA^* may be caused by elimination of sites near FCD1 of FRQ (Figure 1A), a domain required for CK1 interaction and phosphorylation of FRQ N-terminus. *frq^472-536pA^* and *frq^537-570pA^* are 6 and 13 hrs longer respectively than WT, while *frq^472-570pA^* is ~24 hrs longer (Figure 6B), which is significantly longer than 19 hrs (6+13 hrs), suggesting that the additive effect of phosphorylations on period lengths can be stronger than adding up period changes. *frq^472-536pA^* contains three mutations in and close to one of the only two regions of FRQ predicted to have structure (Figure 1B), a region that comprises the FCD2, so the lengthened periods of the two mutants may be due to the reduced CK1 interaction, consistent with an observation that the period length is determined by FRQ-CK1 interaction (Liu *et al*. 2019).

### Epistasis analyses of different phosphogroups on FRQ

An intermediate period length has been observed when different mutations on FRQ were intramolecularly combined, such as *frq^3^* and *frq^7^* (Aronson *et al*. 1994b), *frq^S548A,S900A^* (Baker *et al*. 2009), and *frq^M9+18^* (Tang *et al*. 2009). To test whether phosphorylation in different regions of FRQ have an interplay effect, phosphomutations were combined together. For example, both *frq^S238A, S240A^* and *frq^S238A, S240A, S390A, S392^* display a ~ 2 hrs longer period than WT (Baker *et al*. 2009); the combination of S238A, S240A, S390A, S392, and S394A would be predicted to be ~26 hrs, close to what is seen in *frq^S238A, S240A, S238A, S240A, S390A, S392, S394A^* (27.1 hrs, Figure 7A). Similarly, mutations in *frq^S538A, S540A^* (5 hours longer than WT [+5 hrs]), *frq^S541A, S545A^* (+3 hrs), and *frq^S632A, S643A^* (+3 hrs) (Baker *et al*. 2009) were all introduced to a single *frq* mutant together, *frq^S538A, S540A, S541A, S545A, S632A, S634A^*, which exhibits a rhythm of 32.2 hrs (Figure 7A), exactly what is anticipated from the three original mutants. Taken together, these data indicate an additive effect of certain FRQ phosphomutations on period length.

**Figure 7.**
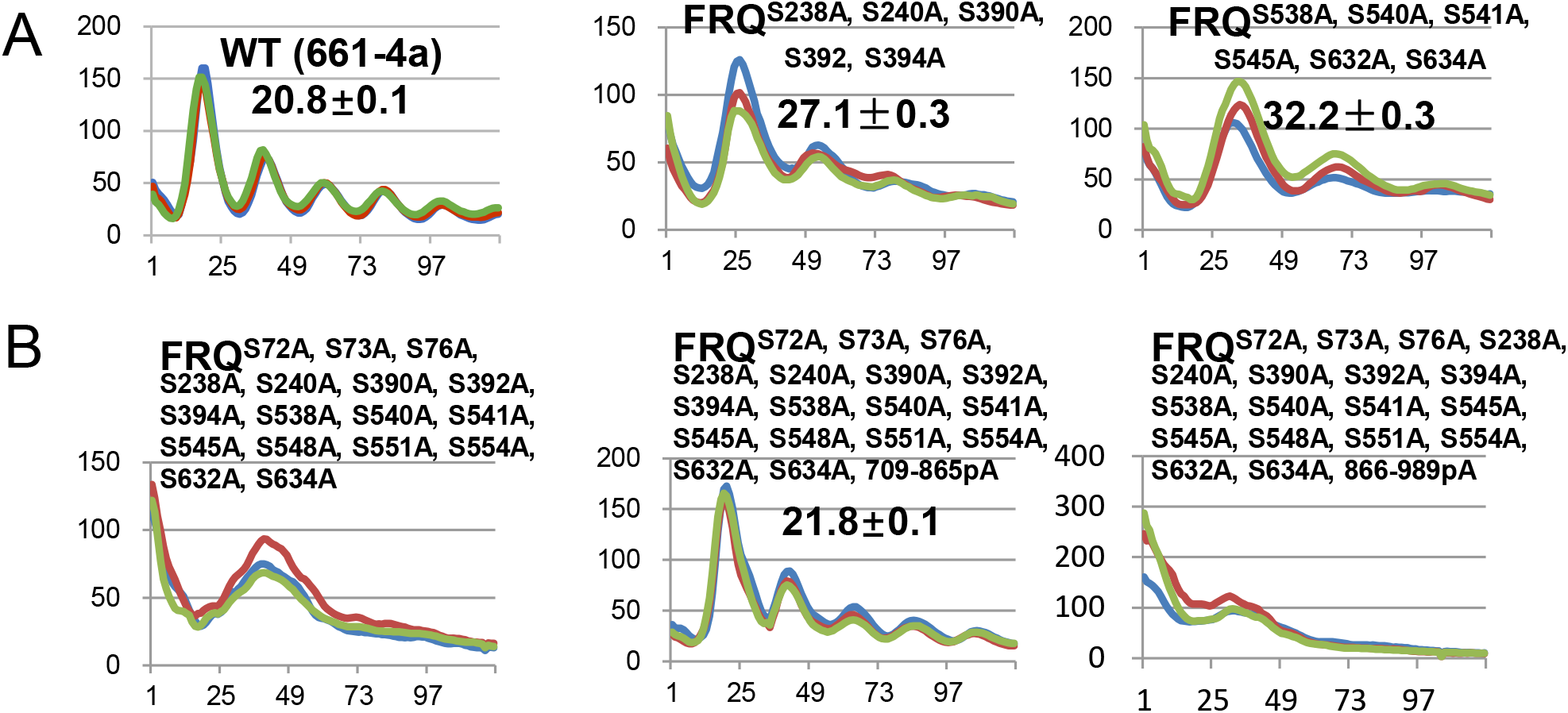
Interplay between FRQ phosphorylations in period determination. **(A)** Luciferase analyses of *frq^S238A, S240A, S390A, S392, S394A^* and *frq^S538A, S540A, S541A, S545A, S632A, S634A^* at 25 °C. (**B**) Luciferase analyses of *frq^S72A, S73A, S76A, S238A, S240A, S390A, S392A, S394A, S538A,S540A, S541A, S545A, S548A, S551A, S554A, S632A, S634A^*, *frq^S72A, S73A, S76A, S238A, S240A, S390A, S392A, S394A, S538A, S540A, S541A, S545A, S548A, S551A, S554A, S632A, S634A, 708-865pA^*, and *frq^S72A, S73A, S76A, S238A, S240A, S390A, S392A, S394A, S538A, S540A, S541A, S545A, S545A, S548A, S551A, S554A, S632A, S634A, 865-989pA^* at 25 °C.

To examine the period lengthening effect caused by individual mutations, FRQ phosphomutations causing increased period lengths (Baker *et al*. 2009) were together introduced into a single *frq* mutant, *frq^S72A, S73A, S76A, S238A, S240A, S390A, S392A, S394A, S538A, S540A, S541A, S545A, S548A, S551A, S554A, S632A, S634A^*, which, unexpectedly, displays a loss of rhythmicity (Figure 7B). When combined with mutations in *frq^709-865pA^* (Figures 3C and 4), *frq^S72A, S73A, S76A, S238A, S240A, S390A, S392A, S394A, S538A, S540A, S541A, S545A, S548A, S551A, S554A, S632A, S634A, 709-865pA^*, surprisingly, fully restores rhythmicity in *frq^S72A, S73A, S76A, S238A, S240A, S390A, S392A, S394A, S538A, S540A, S541A, S545A, S548A, S551A, S554A, S632A, S634A^* with a period length very similar to that in *frq^768-865pA^* (Figure 7B). However, *frq^S72A, S73A, S76A, S238A, S240A, S390A, S392A, S394A, S538A, S540A, S541A, S545A, S548A, S551A, S554A, S632A, S634A, 866-989pA^* still behaves arrhythmically as *frq^S72A, S73A, S76A, S238A, S240A, S390A, S392A, S394A, S538A, S540A, S541A, S545A, S548A, S551A, S554A, S632A, S634A^* (Figure 7B). Consistent with the data above, *frq^709-989pA^* displays a circadian rhythm of ~15 hrs, which is the same as *frq^866-989pA^* rather than *frq^709-865pA^* (Figure 3 B and C), suggesting that the 11 phosphoevents in aa 866-989 are epistatic to the 9 ones found in aa 709-865. Collectively, these data suggest that the interplay between phosphogroups on FRQ can be varied, including an averaging, additive, or epistatic effect on the period length and rhythmicity.

### Beyond its negative charge, the phosphate group at pS900 regulates FRQ activity at the structure level

Phosphomimetics by amino acid substitutions like Asp (D) or Glu (E) are a widely used strategy to simulate phosphorylation, by constitutively introducing a negative charge into a domain. In *Neurospora*, phosphomimetics have been successfully employed to study phosphorylation of the core clock components, WC-1, WC-2 and FRQ at certain sites, such as *wc-1^S971D^* (Wang *et al*. 2019), *wc-2^15pD^* (Wang *et al*. 2019), and *frq^S548D^* (Baker *et al*. 2009), revealing interesting effects caused by constant phosphorylation at these sites. To further examine the mechanism behind the phosphorylation-controlled change of FRQ activity, several key phosphosites on FRQ were mutated to Asp (D) or Glu (E) to mimic the negative charge of the phosphate group. The period lengths of *frq^S915A, S917A^* and *frq^S923A^* are ~2 and 1 hr longer respectively (Baker *et al*. 2009), whereas *frq^S915D, S917D, S923D^* and *frq^S915E, S917E, S923E^* shows a WT period (Figure 8A); similarly, *frq^S548A^* has a 4-hrs period lengthening, while *frq^S548D^* maintains a WT period (Baker *et al*. 2009), suggesting that phosphorylation at these sites of FRQ plays a role in maintaining the pace of the clock. *frq^S900A^* displays a period of ~18 hrs (Baker *et al*. 2009), but, unexpectedly, both *frq^S900D^* and *frq^S900E^* exhibit the same period length (18.4 and 18.9 hrs respectively) as *frq^S900A^* (Figure 8B), suggesting that the structure of the phosphate group of pS900 plays a more important role than the negative charge that it carries in tuning the FRQ activity. Although the phosphate group and Asp/Glu are both negatively charged, they are structurally distinct from each other, which may explain why both *frq^S900D^* and *frq^S900E^* failed to mimic phosphorylation at S900 of FRQ but instead behave like phosphorylation elimination.

**Figure 8.**
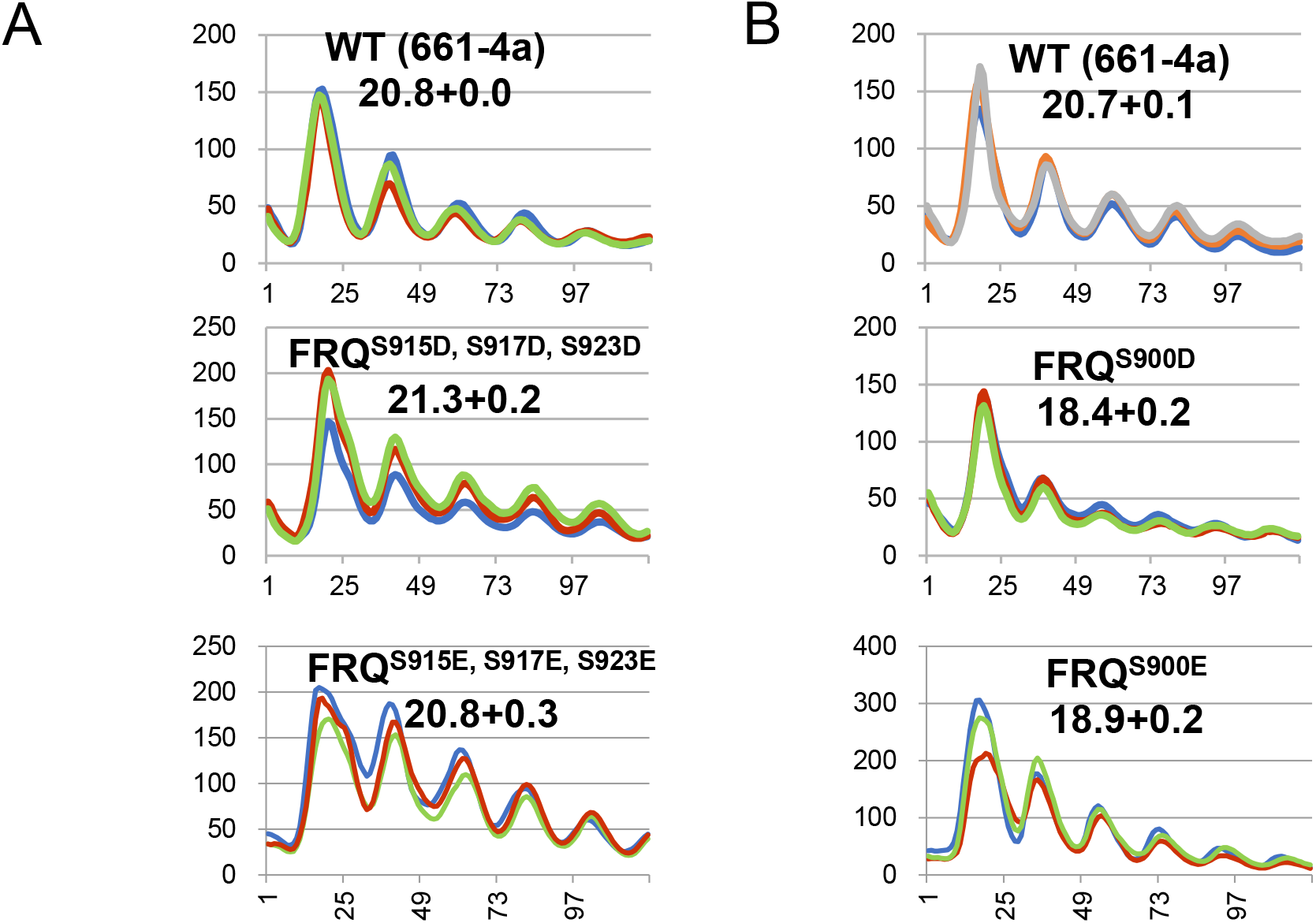
Phosphomimetics of residues on FRQ shows opposite effects. **(A)** *frq^S900D^* and *frq^S900E^* display the same period length as *frq^S900A^* at 25 °C. **(A)** *frq^S915D, S917D, S923D^* and *frq^S915E, S9157E, S923E^* show a WT period length at 25 °C.

## Discussion

FRQ has been predicted to be a largely unstructured protein comprising many disordered regions that make most of its residues exposed and accessible by kinases in the cell (Hurley *et al*. 2013; reviewed in Pelham *et al*. 2020; Marzoll *et al*. 2022b, 2022a), which is consistent with a large number of phosphorylatable residues identified on it. Although over 100 phosphosites have been identified (Baker *et al*. 2009; Tang *et al*. 2009) and partially confirmed by (Horta *et al*. 2019) and Ala mutations to some of these phosphoresidues have been shown to alter period lengths, their functions are still largely unknown due to lack of systematic mutagenetic analyses to all the phosphoevents on FRQ. In this study, we generated and studied *frq* phosphomutants covering all 110 phosphosites, and detailed mutagenetic analyses have allowed circadian roles of these site assigned to different domains of FRQ. Excluding those mutations that resulted in arrhythmicity, we found that mutating phosphoresidues in the N-terminal or middle regions of FRQ only cause increased or unaltered period lengths while removal of phosphorylated residues at the C-terminus results in a decreased period length and an over-compensated circadian clock in a physiological temperature range. Interestingly, either an additive or epistatic effect on rhythmicity has been observed when combining different groups of mutations together.

How is FRQ activity tightly tuned over the course of a day? Recent publications have strongly challenged the model in which the period length is determined by the half-life of FRQ and instead support that time-of-day-specific phosphorylation of FRQ finely controls its activity (Baker *et al*. 2009; Larrondo *et al*. 2015; Liu *et al*. 2019; Hu *et al*. 2021). Lacking an enzymatic activity, FRQ mainly acts as a molecular platform that recruits kinases to phosphorylate its transcription activator, WCC, thereby closing the feedback loop. An intramolecular interaction between the N- and C-termini of FRQ has been demonstrated (Querfurth *et al*. 2011), which might be weakened or disrupted by progressive phosphorylation over time, leading to decreased interaction or dissociation between FRQ and its interactors, removal of the repression on WCC, and restarting the next circadian cycle. FRQ phosphorylation can impact its activity through two different ways: Phosphorylation occurring within or close to a domain(s) can directly alter its function and interacting partners; most phosphosites are located in the disordered regions of FRQ, and they can change the overall structure of FRQ by disrupting the intramolecular interaction between its N- and C-termini, which is essential for the FRQ activity (Querfurth *et al*. 2011). If phosphorylation at N- and middle regions of FRQ is not allowed or occurs at a slow pace, then the intramolecular interaction within FRQ will be sustained longer and so does the capacity of FRQ in WCC repression, consistent with the long periods seen in the *frq* mutants (Figures 3B and 3C). Phosphorylation of the FRQ C-terminal tail plays a role in slowing down the pace of the feedback loop; if this molecular brake via phosphorylation is broken, FRQ loses its capacity of promoting WCC phosphorylation more quickly so WCC regains its transcription activity sooner, reflected by the short periods seen in mutants, such as *frq^709-989pA^* and *frq^866-989pA^* (Figures 3B and 3C). High temperatures might be able to compensate for the loss of phosphorylation in the C-terminal tail of FRQ, so the shortened period gets rescued to some extent at a higher temperature (Figure 4).

Both FRQ and its transcription activator WCC are subject to extensive phosphorylations in a circadian cycle, and similarly, activities of both protein complexes are finely controlled by multiple phosphoevents (Baker *et al*. 2009; Tang *et al*. 2009; Wang *et al*. 2019). For example, WCC transcription activity in the dark is completely inhibited only when a small group of sites on both WC-1 and WC-2 are simultaneously phosphorylated (Wang *et al*. 2019) while a large number of phosphoevents on WCC play no or little role in the core clock but only act on lowering expression of *frq* and clock-controlled genes (namely circadian amplitude) (Wang *et al*. 2019). Similarly, although FRQ is also heavily phosphorylated at multiple sites over time, to date no single phosphomutant of *frq* has been found to be constantly active or inactive in a circadian cycle, suggesting that FRQ activity is determined by multiple phosphoevents. However, an obvious difference between phosphorylation on FRQ and WCC is that most *wcc* phosphomutants do not show an altered period length (Wang *et al*. 2019), whereas a large number of *frq* phosphomutants spanning the whole protein display period changes to a certain extent (Mehra *et al*. 2009; Baker *et al*. 2009; Tang *et al*. 2009; Larrondo *et al*. 2015). These observations are consistent with a model in which complexing with FRH, FRQ serves as a platform by recruiting kinases to phosphorylate and inhibit WCC, so multiple domains of FRQ participate in recruiting many other proteins, including FRH, CKI, and FRQ itself via FFD, FCD, and CC domains (Figure 1B), respectively, as well as multiple regions for interaction with WCC (data not shown), and correspondingly phosphorylations near or within these regions may directly or indirectly regulate these interactions; FRQ-dependent repression on WCC mainly targets the DNA binding domain and its nearby regions of WCC (Wang *et al*. 2016, 2019), which explains why mutations to phosphosites in other regions of WCC do not dramatically impact the period length.

FRQ phosphorylation dynamics have been investigated by quantitative mass spectrometric analyses including stable isotope labeling by amino acid in cell culture (SILAC) (Baker *et al*. 2009) and N^15^/N^14^ isotope labelling (Tang *et al*. 2009). A cluster of residues surrounding the PEST-2 region (near aa 795–929) become hyperphosphorylated at CT8 when the level of new FRQ and FRQ activity start to increase. Eliminating phosphorylation in 709-989 (*frq^709-989pA^*) results in a short period (Figure 3B), suggesting phosphorylation in this region may be required for FRQ activity in repressing WCC. Sites specific to the N-terminus of l-FRQ become phosphorylated at CT16, a late time point; sites in the PEST-1 domain (aa 537–558) become hyperphosphorylated later, peaking at CT12, suggesting that these phosphorylations may function in inhibiting FRQ activity. Consistent with these, *frq^1-114pA^* and *frq^536-570pA^* develop long periods of 26.7 and 34.8 hrs, respectively (Figures 3C and 5B). Phosphorylation of aa 211–257 peaks earlier and decreases relatively over time, suggesting that the dynamics of phosphorylation at these regions correlates with and may impact the change of FRQ activity in a circadian cycle (Baker *et al*. 2009), consistent with the arrhythmicity seen in *frq^211-257pA^* (Figure 6A). Due to scarcity of purified FRQ for in vitro studies and potential ionization issues of peptides bearing multisite phosphorylations in mass spectrometry, how phosphorylation of FRQ at many sites may change in concert in a circadian cycle is still largely unknown, which restricts our understanding of the role of time-specific phosphoclusters on FRQ.

Results in this work may inform understanding of mammalian and insect clocks, many facets of which are also built on time-specific multisite phosphorylation events to the key components (reviewed in (Brenna and Albrecht 2020)). PER/TIM in *Drosophila* and PERs/CRYs in mammals act as the negative elements in the negative feedback loop by inhibiting Clk/Cyc and CLOCK/BMAL1 activities respectively, terminating their own expression and thereby closing the circadian negative feedback loop. Similar to FRQ and WCC in *Neurospora*, PER/TIM and PERs/CRYs also undergo extensive phosphorylation, and phosphorylation of PER/TIM and PERs/CRYs has been shown to be a critical mechanism in controlling both the fly and mammalian clocks (Chiu *et al*. 2008, 2011; Lamia *et al*. 2009; Top *et al*. 2016; Cao *et al*. 2021; Cai *et al*. 2021; An *et al*. 2022). The strategy adopted here to progressively dissect scores of phosphosites on FRQ might be applicable to facilitating identification of essential phosphoevents on clock components in other systems.

## Acknowledgements

This work was supported by a grant from the National Institutes of Health to J.C.D. (R35GM118021).

## Materials and Methods

### Growth conditions

All vegetative cultures were maintained on complete medium slants bearing 1 x Vogel’s, 1.6% glycerol, 0.025% casein hydrolysate, 0.5% yeast extract, 0.5% malt extract, and 1.5% agar (Vogel 1956). Sexual crosses were performed on Westergaard’s agar plates containing 1 x Westergaard’s salts, 2% sucrose, 50 ng/mL biotin, and 1.5% agar (Westergaard and Mitchell 1947).

### *frq* mutant generation

To lower the cost of making a large number of *frq* mutants, a method described in (Baker *et al*. 2009) was modified to use yeast homologous recombination-based integration of PCR fragments (Wang *et al*. 2014) bearing FRQ point mutations to restriction-digested *pCB05* in place of the QuickChange II® Site-directed Mutagenesis Kit (Stratagene). Four primer sets were used as flanks to facilitate homologous recombination in a yeast strain (*FY834*) by which point mutations of *frq* were introduced from PCR primers. To introduce mutations to aa 1-214 of FRQ, two PCR reactions were performed: one with a forward primer “*frq* segment 1F” (5’-GAACCAGAACGTAGCAGTGTG-3’) and a reverse primer “#pA R” bearing a point mutation(s) to FRQ and the other using a forward primer “#pA F” which is reverse and complementary to “#pA R” and a reverse primer “*frq* segment 1R” (5’-GACGATGACGACGAATCGTG-3’), and then the two PCR products were cotransformed into yeast along with *pCB05* digested with *Bst*XI and *Xho*I to create a circular construct. Similarly, to introduce mutations falling in aa 215-437 of FRQ, primers “*frq* segment 2F” (5’-GTGAGTTGGAGGCAACGCTC-3’) and “*frq* segment 2R” (5’-GTCCATATTCTCGGATGGTA-3’ were used for PCRs in combination with *pCB05* digested with *Xho*I to *Nru*I; “*frq* segment 3F” (5’-GTCGCACTGGTAACAACACCTC-3’) and “*frq* segment 3R” (5’-CAGCACATGT TCAACTTCAT CAC-3’) were designed for *pCB05* digested with *Nru*I and *Fse*I (FRQ aa 438-675), and “*frq* segment 4F” (5’-CACCGATCTTTCAGGAGACCCTG-3’) and “*frq* segment 4R” (5’-CACTCAGGTC TCAATGGTGA TG-3’) work for *pCB05* digested with *Fse*I *and Mlu*I (FRQ aa 676-989).

If multiple phosphosites span two or more PCR segments above, corresponding restriction enzymes and primers encompassing the region were chosen and combined for recombination in yeast. All mutations were verified by cycle sequencing at the Dartmouth Core facility. The open reading frame of *frq* bearing 84 phosphomutations (*frq^84A^*) from (Baker *et al*. 2009) was custom-synthesized and purchased from Genscript, and to *frq^84A^*, additional 26 phosphosites identified in (Tang *et al*. 2009) were further mutated to Ala by PCR reactions using primer pairs bearing mutations to create *frq^110pA^*. All *frq* mutant constructs were targeted by homologous recombination to its native locus. Plasmids verified by cycling sequencing were linearized with *Ase*I and *Ssp*I and PCR-purified for *Neurospora* transformation. *Neurospora* transformation was performed as previously reported (Colot *et al*. 2006). The recipient strain used in transforming *frq* mutants is Δ*frq::hph; Δmus-52::hph; ras-1^bd^; C-box luc at his-3*, and all *frq* mutants made in this study were in the *ras-1^bd^* genetic background (Belden *et al*. 2007) and bear a V5H6 tag at their C termini and *frq C-box*-driven codon-optimized firefly *luciferase* gene at the *his-3* locus (Gooch *et al*. 2008), except for the strains in Figure 5, all of which bear *frq-C-box* driven *luciferase* at the *csr-1* locus rather than *his-3*. These strains were constructed by crossing phosphomutants from Baker *et al*. (2009) to *frq-C-box-luc*.

### Immunoprecipitation (IP)

IP was performed as previously described (Wang *et al*. 2016). Briefly, 2 mgs of total protein were incubated with 20 μL of V5 agarose (Sigma-Aldrich, #7345) as indicated rotating at 4 °C for 2 hrs. The agarose beads were then washed twice with the protein extraction buffer (50 mM HEPES [pH 7.4], 137 mM NaCl, 10% glycerol, 0.4% NP-40) and eluted with 100 μL of 5 × SDS sample buffer heated at 99 °C for 5 min.

### Lambda protein phosphatase-treatment of FRQ

V5H6-tagged FRQ was immunoprecipitated with 20 μL of V5 agarose (Sigma-Aldrich, Catalog #7345) from 2 mg of centrifugation-cleared lysate, FRQ-bound V5 agarose was thoroughly washed twice using the protein extraction buffer, and all supernatant was carefully removed by pipetting. To make a total reaction volume of 50 μL, 40 μL of H_2_O, 5 μL of 10 x NEBuffer for Protein MetalloPhosphatases (PMP), 5 μL of 10 mM MnCl_2_, and 2 μL of lambda protein phosphatase (NEB, Catalog #P0753S) were added to the washed FRQ-coupled V5 resin. The mixture was incubated at 30 °C for 30 minutes, and then 50 μL of 5 × SDS sample buffer was added and heated at 99 °C for 5 min (Zhou *et al*. 2018).

### Western blot (WB)

For WB, equal amounts (15 μg) of cleared protein lysate were loaded per lane in an SDS-PAGE gel. FRQ, FRH, WC-1, and WC-2 antibodies were previously described (Garceau *et al*. 1997; Denault *et al*. 2001; Froehlich *et al*. 2002). Antibody against V5 (Thermo Pierce) was used at 1:5,000 dilution as the first antibody in WB (Wang *et al*. 2021).

### Phos-tag gel

To better resolve FRQ phosphorylation events, Phos-tag chemical purchased from ApexBio was added at the final concentration of 20 μM to the 6.5% SDS-PAGE Tris-Glycine gel with a ratio of 149:1 acrylamide/bisacrylamide (Wang *et al*. 2019).

### Luciferase assay

Luciferase assays were performed as previously described (Larrondo *et al*. 2012). 96-well plates with each well containing 0.8 mL of the luciferase assay medium were inoculated with conidial suspension and unless otherwise specified, strains in luciferase assays were cultured at 25 °C and in constant light for 16–24 hrs and then transferred to the dark at the same temperature for recording light signals. Bioluminescence signals were recorded with a CCD camera every hour, data were obtained with ImageJ and a custom macro, and period lengths of the strains were manually calculated. Raw data from three replicates are shown, and time (in hours) is on the x-axis while arbitrary units of the signal intensity is on the y-axis. In Figure 4, the strains were synchronized at 20, 25, or 30 °C plus light overnight and then transferred to darkness at the same temperature used in synchronization to monitor light production by a CCD camera. Strains in Figure 5 were entrained at 25 °C for two days on a 12/12 light dark cycle before transferring to the dark at either 20, 25, or 30 °C to monitor light production by a CCD camera. Luciferase assay medium contains 1 x Vogel’s salts, 0.17% arginine, 1.5% bacto-agar, 50 ng/mL biotin, and 0.1% glucose, and liquid culture medium (LCM) contains 1 x Vogel’s, 0.5% arginine, 50 ng/mL biotin and 2% glucose. WT used in luciferase assays was 661-4a (*ras-1^bd^, A*) that contains the *frq C-box* fused to the codon-optimized firefly *luciferase* gene (transcriptional fusion) at the *his-3* locus.

---

**Supporting Figure 1.**
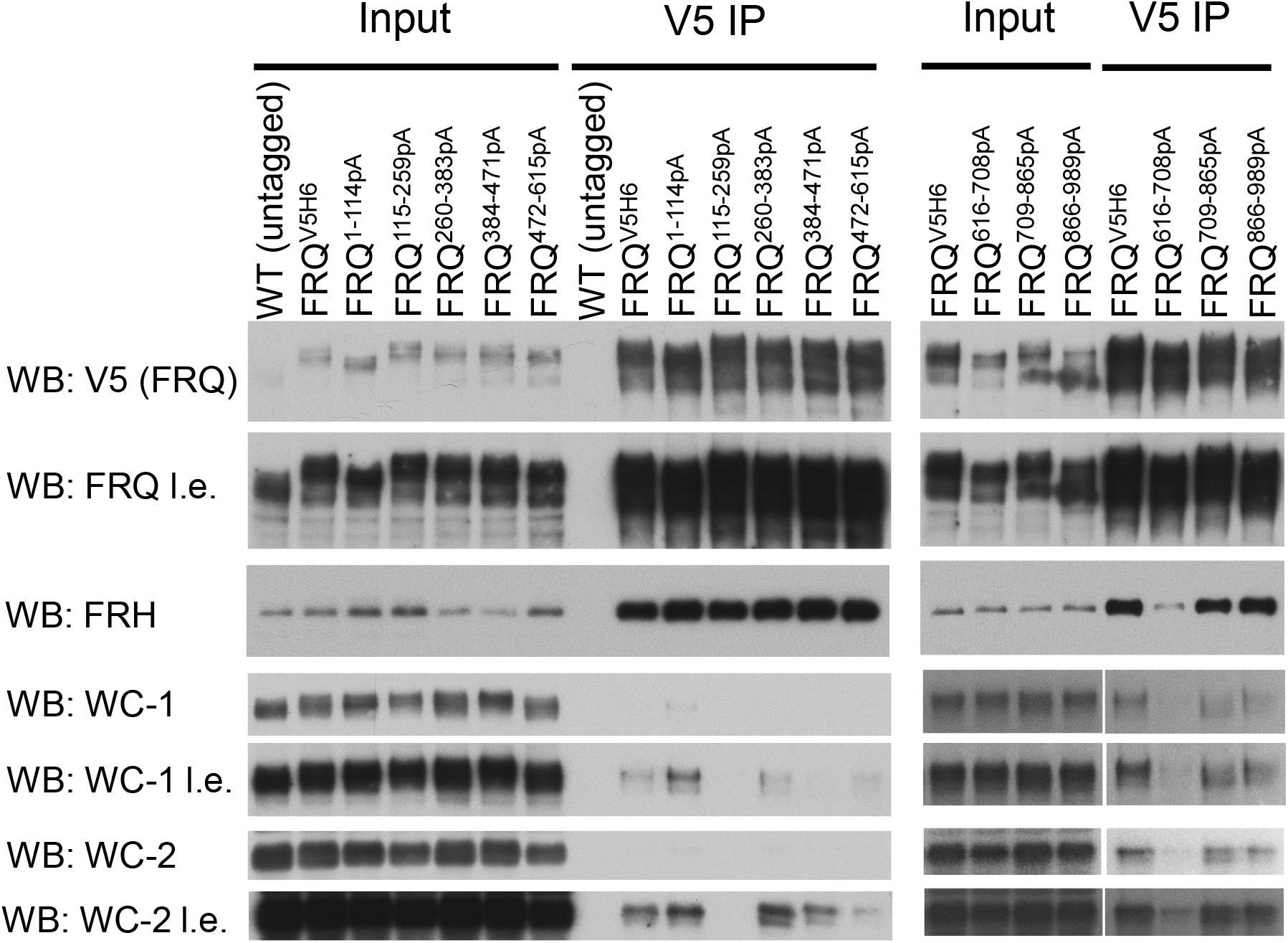
Expression of FRQ, FRH, WC-1, and WC-2 and their interactions in *frq^1-114pA^, frq^115-259pA^, frq^260-383pA^, frq^384-471pA^, frq^472-615pA^, frq^615-708pA^, frq^708-865pA^*, and *frq^865-989pA^* by immunoprecipitation. Strains were cultured at 25 °C in the light, and V5 IP was performed using centrifugation-cleared lysate. Except for WT untagged, FRQ in all other strains bear a V5H6 tag at the C-terminus for detection and immunoprecipitation.

**Supporting Figure 2.**
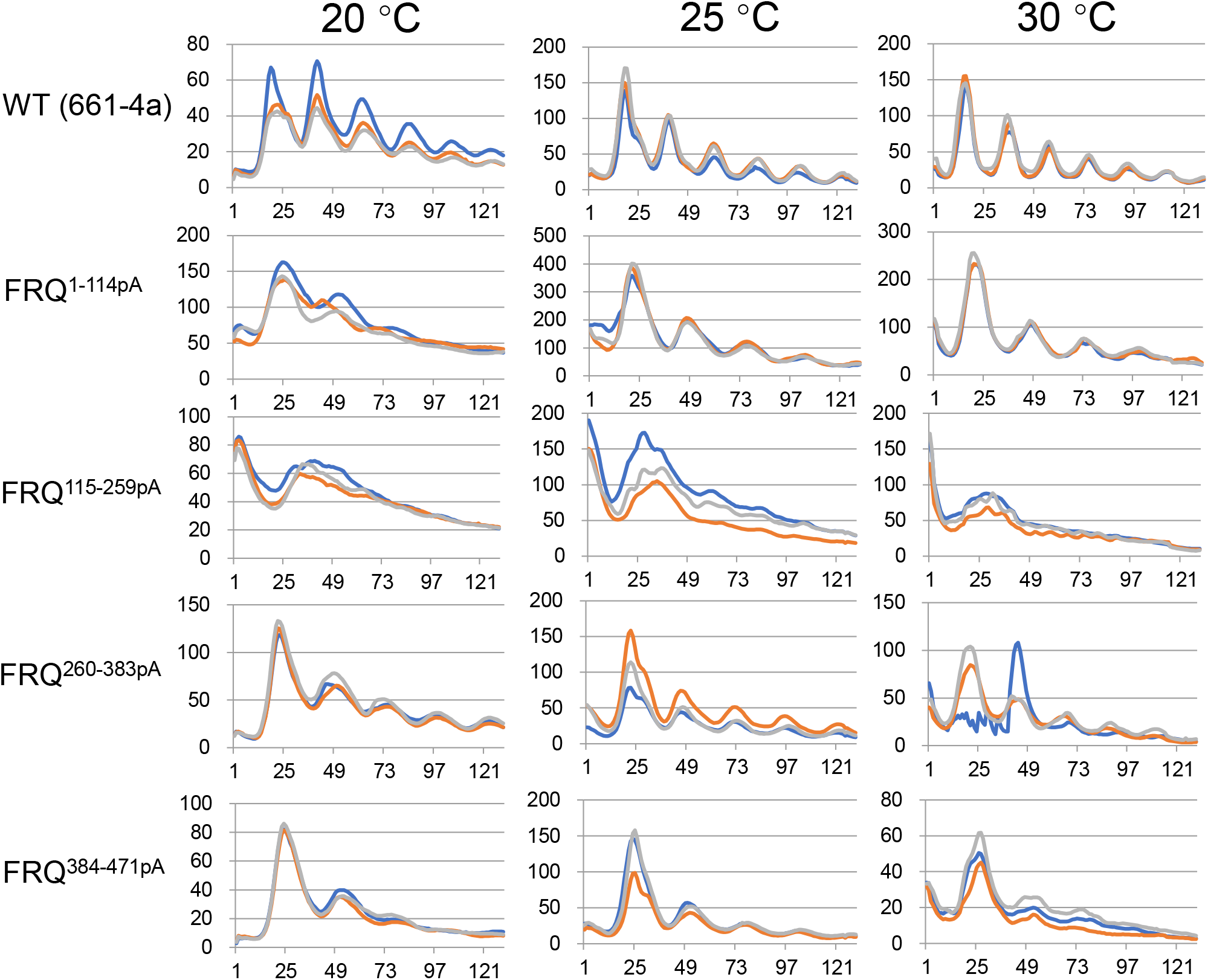
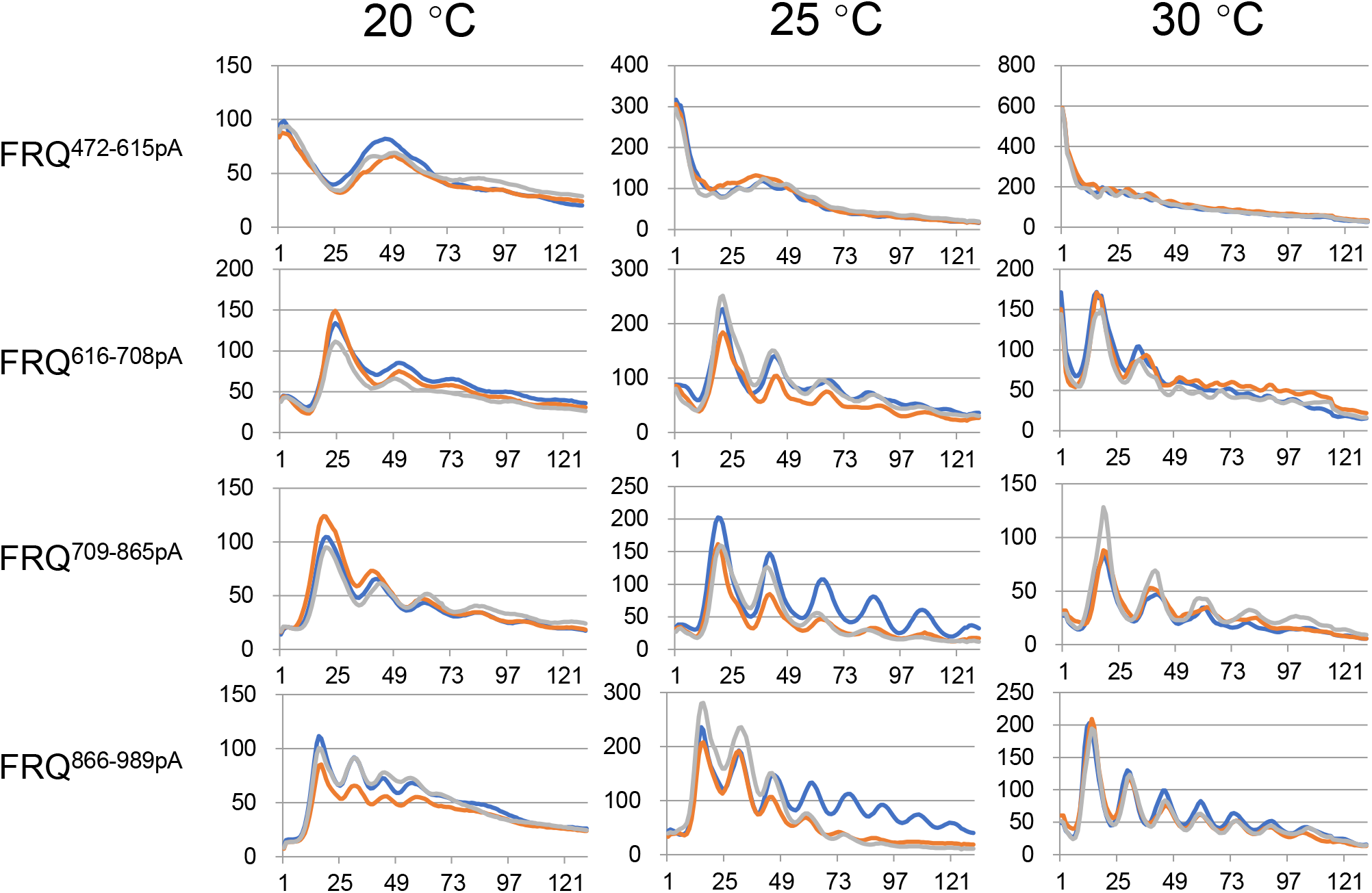
Raw luciferase data of *frq^1-114pA^, frq^115-259pA^, frq^260-383pA^*, *frq^384-471pA^, frq^472-615pA^, frq^616-708pA^, frq^709-865pA^*, and *frq^866-989pA^* at 20, 25, and 30 °C from three replicates. These strains were synchronized at 20, 25, or 30 °C in the light overnight and in the following day transferred to darkness at the same temperature used in synchronization. Luciferase signals were immediately recorded by a CCD camera after transfer of the strains to the dark.

**Supporting Figure 3.**
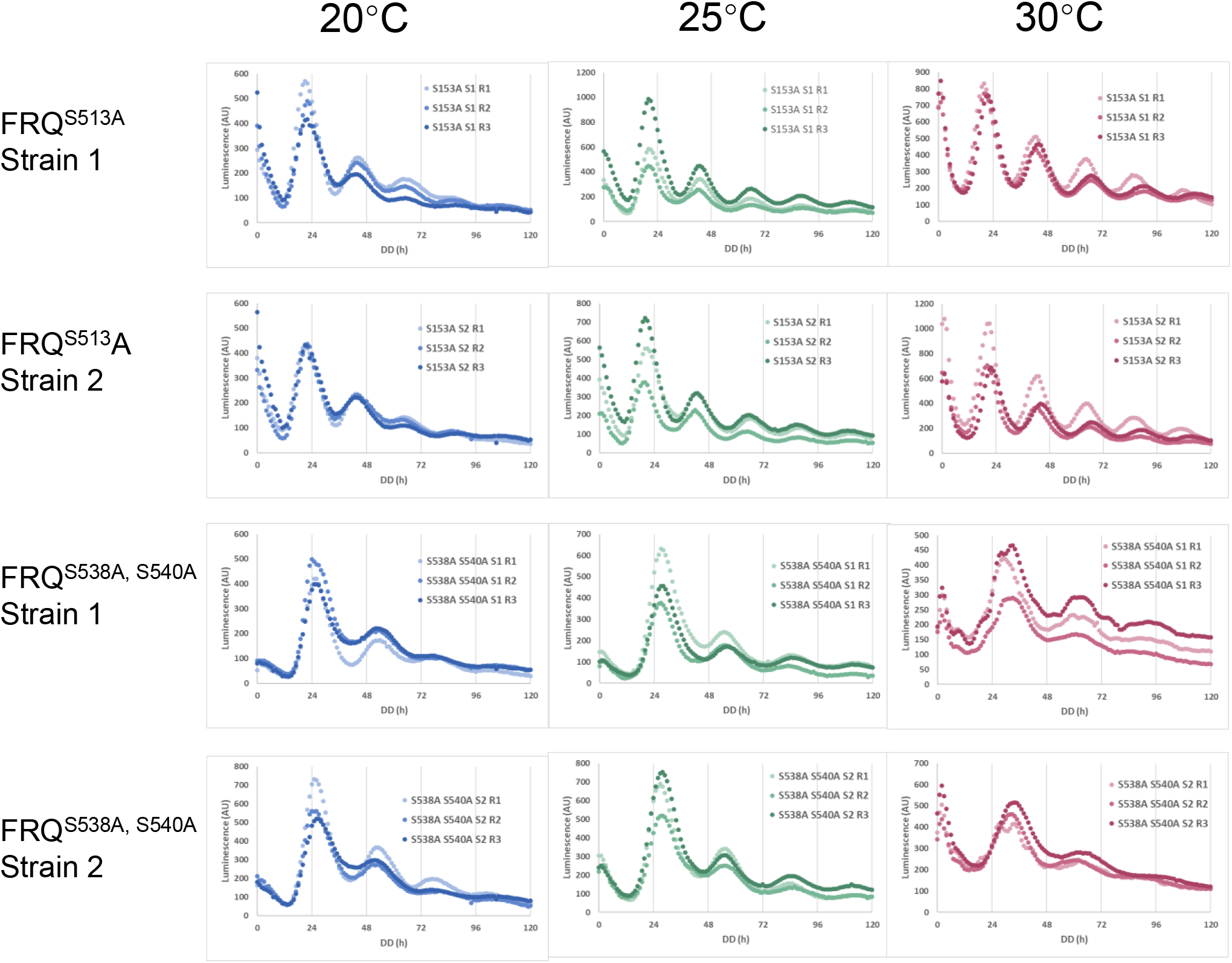
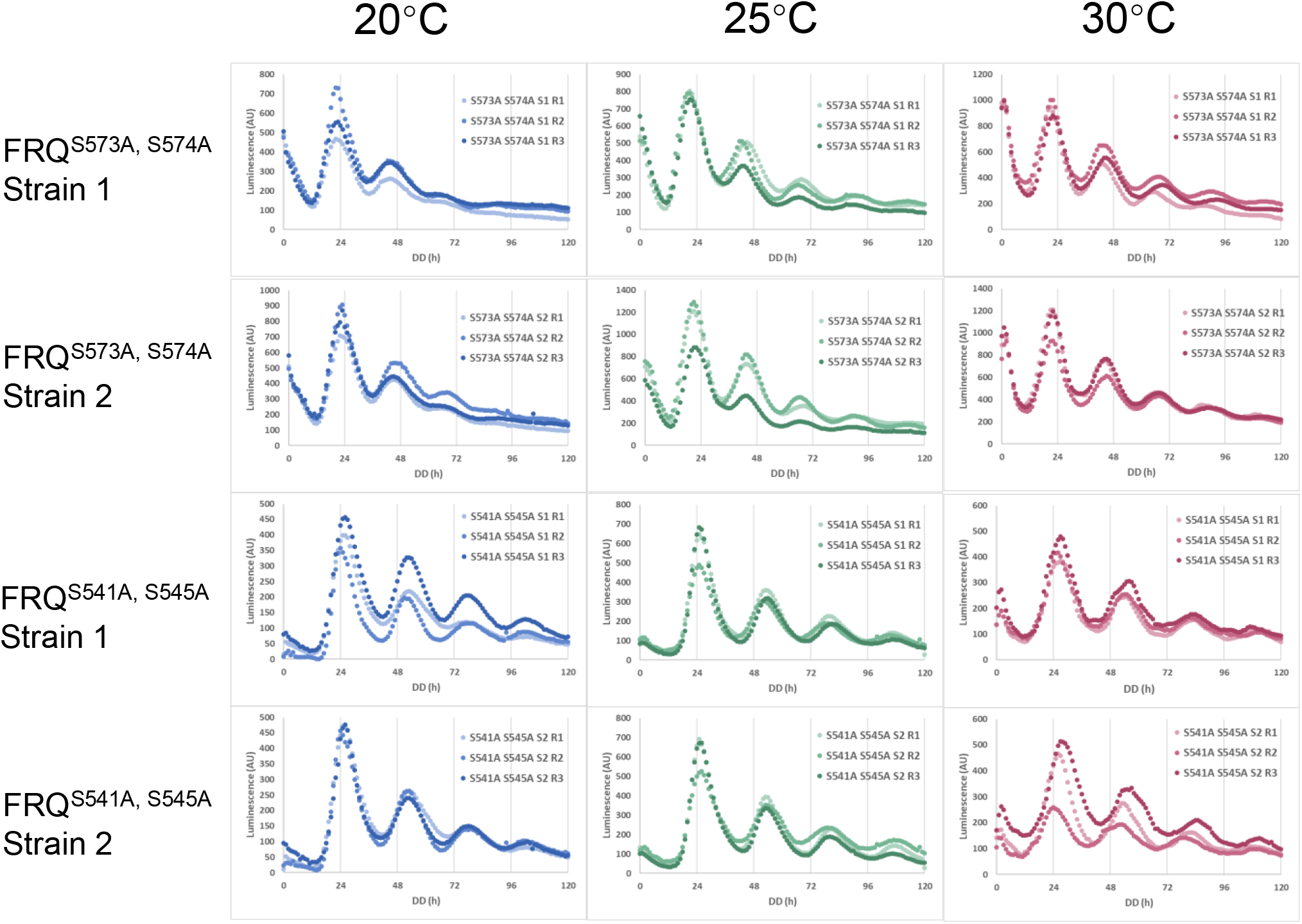
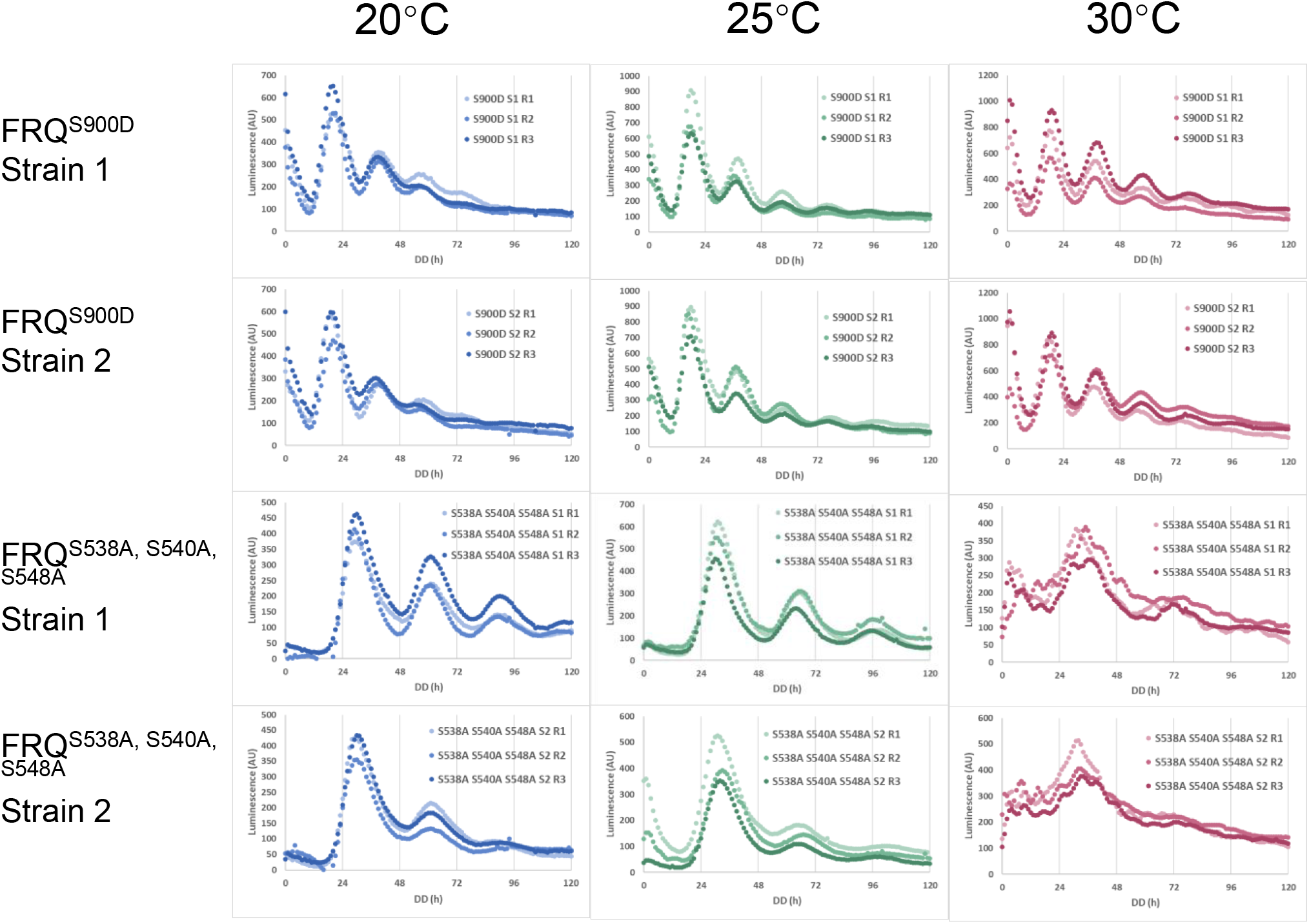
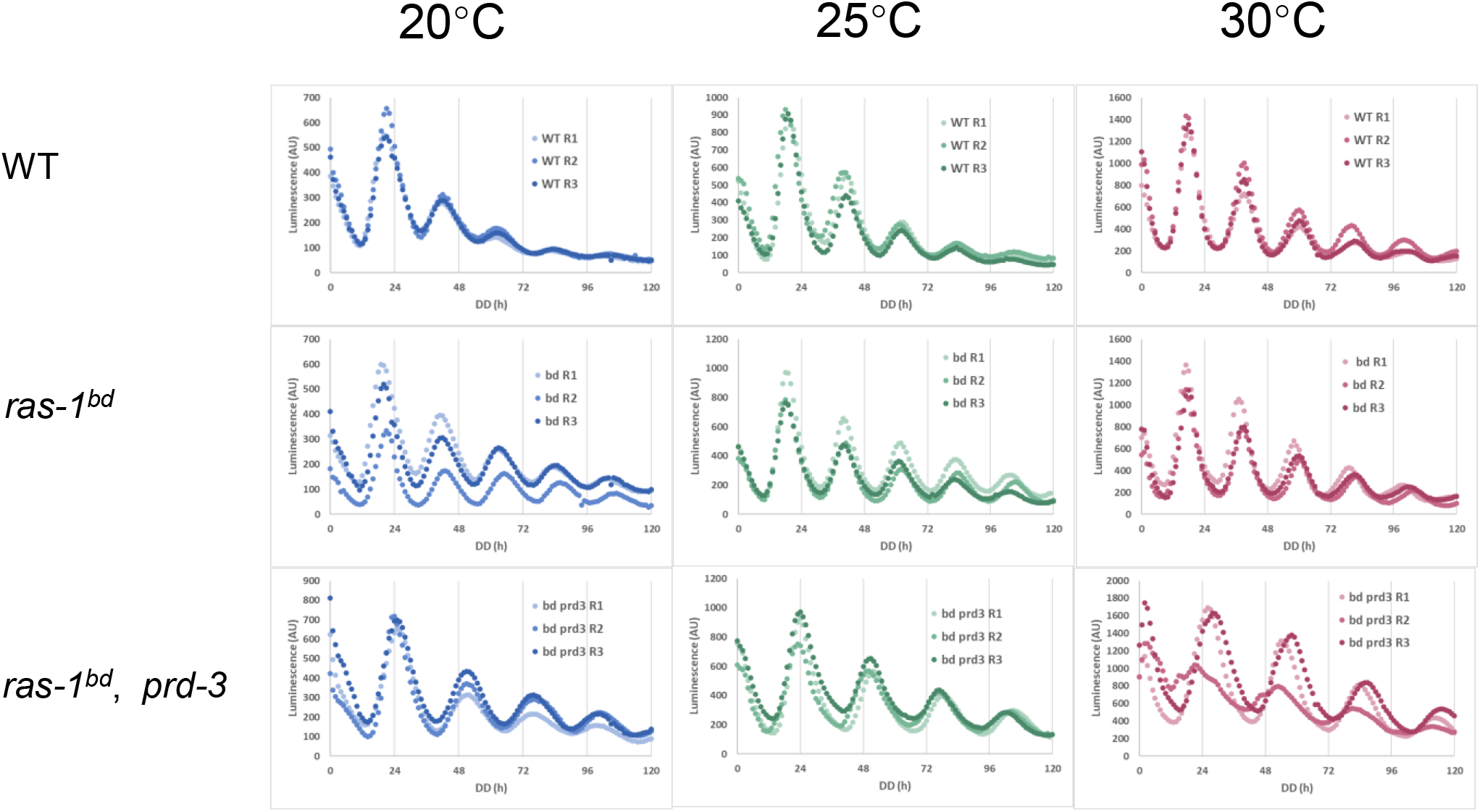
Raw luciferase data of strains in Figure 5 at 20, 25, and 30 °C from three replicates.

**Supporting Table 1** Raw period data and statistics of luciferase analyses in Figure 5.

## Notes

### Competing Interest Statement

The authors have declared no competing interest.

